# Targeting Osteosarcoma heterogeneity to improve therapeutic response

**DOI:** 10.1101/2025.01.02.631118

**Authors:** Brenda Melano, Truc Dinh, Bahar Zirak, Brian J Woo, Benedict Choi, Mehran Karimzadeh, Hani Goodarzi, E. Alejandro Sweet-Cordero

## Abstract

Intra tumor heterogeneity complicates cancer therapy by providing tumors with the ability to alter their phenotypes and become more therapy resistant. Here, we tested the hypothesis that identifying and modulating expression of key state-specific transcription factors could be used as a strategy for driving cells to a more therapy-sensitive state. Recent single-cell studies have explored the inter and intra tumoral heterogeneity of osteosarcoma and identified gene pathways enriched in specific cell states. For example, metastatic tumors are characterized by an expression of genes in the TNF-α, PI3K, TGFß and mTOR pathways. We identified similar profiles in osteosarcoma patient-derived xenograft-derived cell lines and potential transcription factor drivers of these states. We then used perturb-seq to downregulate expression of key transcription factors and evaluated the effect of these modulations on single cell RNA profiles and drug responses. Knockdown of *NFE2L3* or *NR0B1* increased the proportion of cells sensitive to targeted therapy. This approach, which could potentially be applied to other cancers, could be used as a strategy to increase the response to targeted therapies by increasing the proportion of cells in a drug-sensitive state.

**Highlights:**

- Distinct transcriptomic states were identified in osteosarcoma cell lines using single-cell RNA sequencing.
- Lineage tracing identified states with differential sensitivity to therapy.
- Using perturb-seq, we identified transcription factors that drive cells towards a more sensitive state.
- The transcription factor NFE2L3 was identified as targets capable of reprogramming cells to a sensitive state.

## Introduction

Intra-tumor heterogeneity and cell plasticity pose major challenges to successfully targeting cancer.^1,2^ Intra-tumor heterogeneity, which can be inferred from the existence of distinct transcriptional states within a single tumor, results in varying phenotypic responses to therapy. This phenomenon allows some cancer cells to adapt or survive upon treatment, leading to relapse and progression despite initial treatment responses. Cellular plasticity is another therapeutic hurdle as it provides cells the ability to rapidly adapt in response to treatment. Osteosarcoma (OS), a bone cancer most commonly found in children and young adults, exemplifies these challenges, particularly in cases involving metastatic disease.^3,4^

Targeted therapies for OS have so far proven mostly ineffective, particularly for patients who present with metastasis.^5^ Highly toxic chemotherapy, specifically Cisplatin, Doxorubicin and Methotrexate, has been the standard of care for over 40 years.^6,7^ Targeting TFs that regulate cellular heterogeneity may be a potential approach for modulating multiple molecular pathways regulating a resistant phenotype in OS and other cancers.^2,8,9^ Recent studies have explored the complexity of intratumoral OS heterogeneity using single-cell sequencing and cell-state transition modeling, identifying states and pathways associated with relapse and metastasis.^10,11^ For example, OS metastases are enriched for pathways including TGF-ß, VEGFA, PI3K/AKT and mTOR.^11,12^ However, it remains unclear how modulating these pathways changes state proportion of cells in different states, which is critical for addressing compensatory mechanisms.

In this study, we show similarities between cell states within PDX-derived OS cell lines and published single-cell RNA-seq data of primary samples. For example, both the PI3K/AKT/mTOR and VEGFA pathways are enriched in a subset of cells in PDX-derived cell lines. While inhibiting these pathways alone have been found to improve therapeutic efficacy,^13,14^ we sought to identify key transcriptional drivers of these cell states, in order to mitigate the compensatory effects of specific pathway inhibition. To this end, we first identified therapy-sensitive cell-states using lineage tracing (LT). After identifying a sensitive transcriptomic state in OS PDX-derived cell lines, we tested whether perturbing lineage drivers modulated the proportion of cells in the sensitive state. Inhibiting expression of *NFE2L3* or *NR0B1* increased the proportion of cells in a therapeutically sensitive state in OS PDX-derived cell lines, providing proof-of-principle for this approach.

## Results

### Osteosarcoma PDX-derived cell lines are characterized by three states

PDX samples have transcriptional profiles similar to those of the primary tumors from which they were derived.^15^ We previously developed a panel of OS PDX-derived cell lines from osteosarcoma PDXs that have been well characterized with regards to genomic and phenotypic features. The bulk transcriptomes of these lines are highly correlated with that of the primary tumor.^16^ We used three of these cell lines (OS152, OS384 and OS742) derived from metastatic, resected and diagnostic samples respectively to study intra sample heterogeneity as a therapeutic hurdle in OS. OS152 and OS742 both have amplifications of *MYC* while OS384 has a *VEGFA* amplification **(Figure S1A)**. Both *MYC* and *VEGFA* amplifications have been previously associated with metastasis and relapse in OS.^17,18^

These three cell lines were analyzed using single-cell RNA sequencing (scRNA-seq).^19^ 6,490 (OS152), 3,244 (OS384), and 4,766 (OS742) single-cell transcriptomes were successfully generated for these cell lines **(Figure S1B and Methods)**. Using leiden clustering, we identified 5 clusters (populations of transcriptionally similar cells) for each cell line **(Figure 2A)**. We then reduced these initial clusters to 3 three groups using non-negative matrix factorization to decompose the data into 3 components **(Figure 1A, 2B-C and Methods)**. Differential gene expression analysis (DGEA) on the groups, which were assigned state names “A”, “B” and “C”, revealed state-specific gene markers. For example, *COL1A1* and *S100A6* were identified as markers for State “A”, *CENPF* and *UBE2C* for State “B” and *FTH1* and *PSMB6* for State “C” **(Figure 2D)**. State specific markers were then filtered to keep those that appeared in two or more of the samples **(Table 1)**.

**Figure 1:**
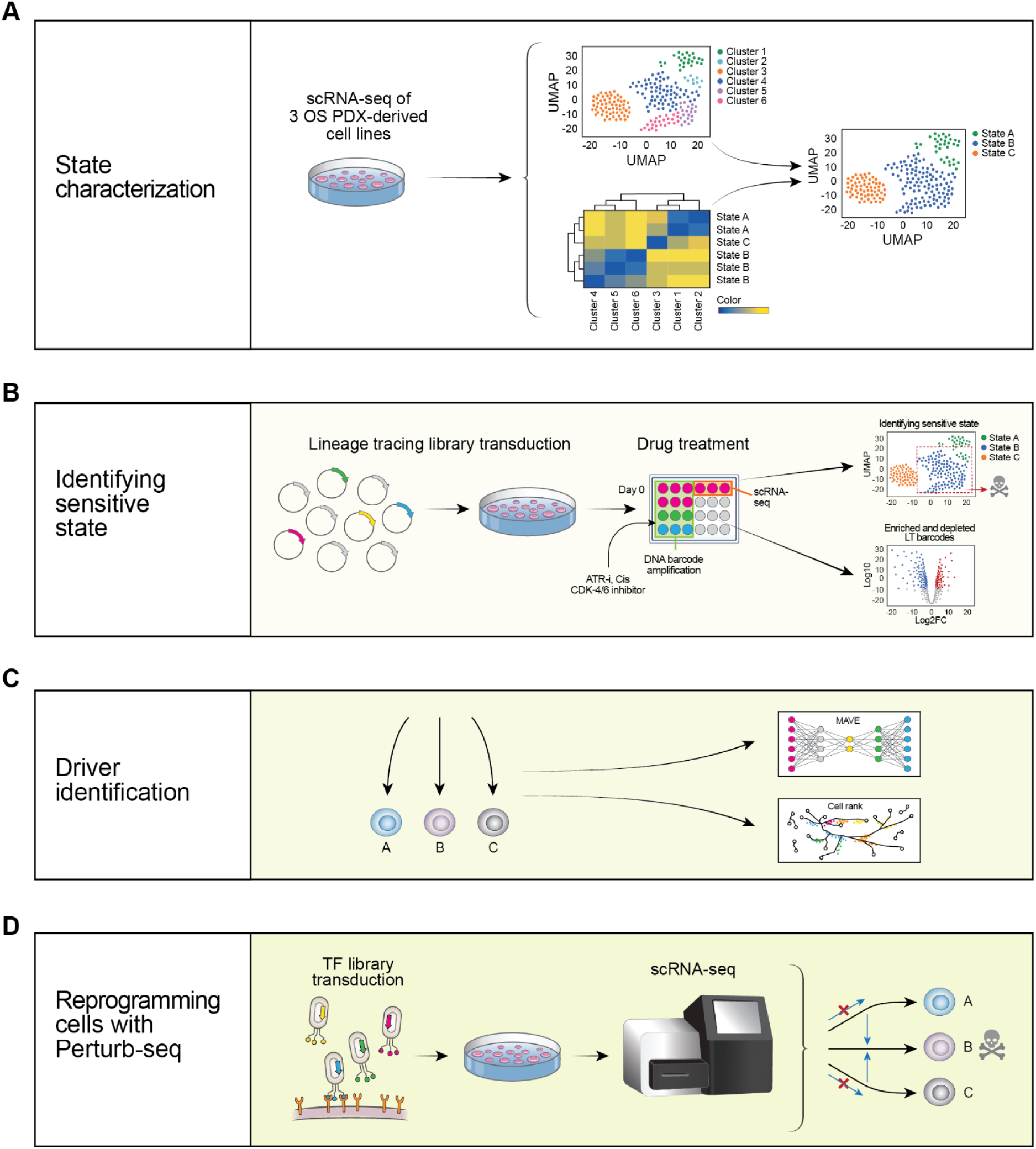
Schematic for target identification pipeline **(A)** Single cell RNA sequencing on the OS PDX-derived cell lines allowed for the aggregation of clusters into three unique transcriptomic states. **(B)** Lineage tracing constructs were transduced into the OS PDX-derived cell lines to test variable clonal sensitivity across various treatment conditions. After isolating the lineage tracing barcodes from both the genomic DNA and scRNAseq libraries, we could identify the sensitive state within the cell lines. **(C)** Candidate State drivers were identified using CellRank and CiberATAC, which uses a biologically informed VAE to identify suspected regulators. **(D)** The perturb-seq pipeline was used to identify which candidate TF regulators could successfully shift cells to the sensitive state.

**Figure 2:**
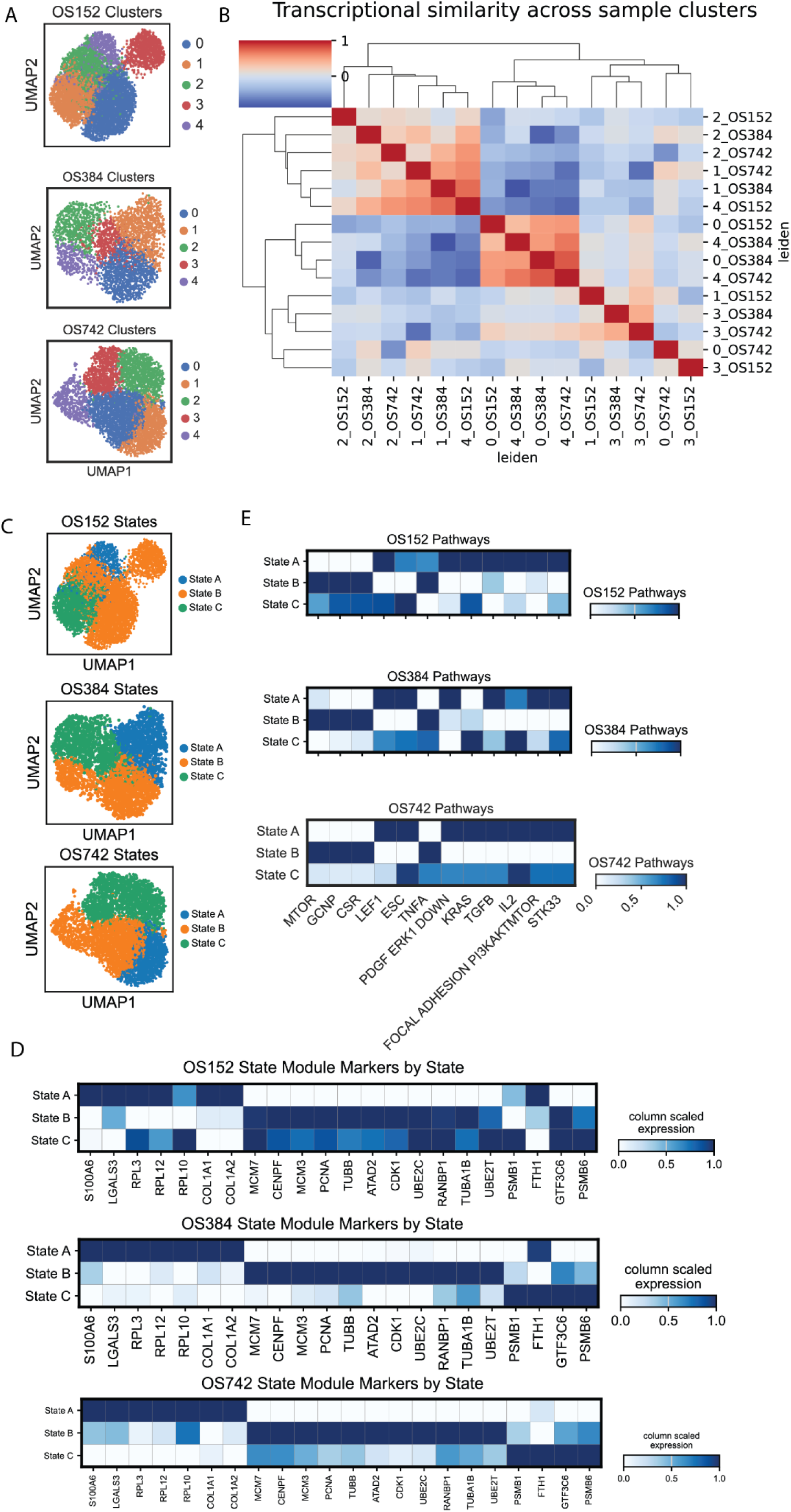
OS PDX-derived cell lines display transcriptomic intra-tumor heterogeneity and subtype classification. **(A)** Uniform manifold approximation and projection (UMAP) of three OS patient derived xenograft (PDX)-derived cell lines: OS152, OS384 and OS742. **(B)** Hierarchical clustering based on cluster-level correlation of mean expression profiles. Red shades represent higher levels of transcriptional similarity. **(C)** Uniform manifold approximation and projection (UMAP) of three OS patient derived xenograft (PDX)-derived cell lines: OS152, OS384 and OS742. These plots show the State “A”, “B” and “C” populations within these cell lines **(D)** Matrix plot illustrating the expression of state module markers across OS152, OS384, and OS742 states. The gene markers are sorted by their state. This plot highlights the gene modules contributing to cellular states. The color bar represents column-scaled expression values. **(E)** To confirm transcriptional similarity on the pathway level and identify which pathways are enriched in the different states, we quantified pathway enrichment in the different states. This analysis was performed on n=3 biological replicates.

**Table 1.**
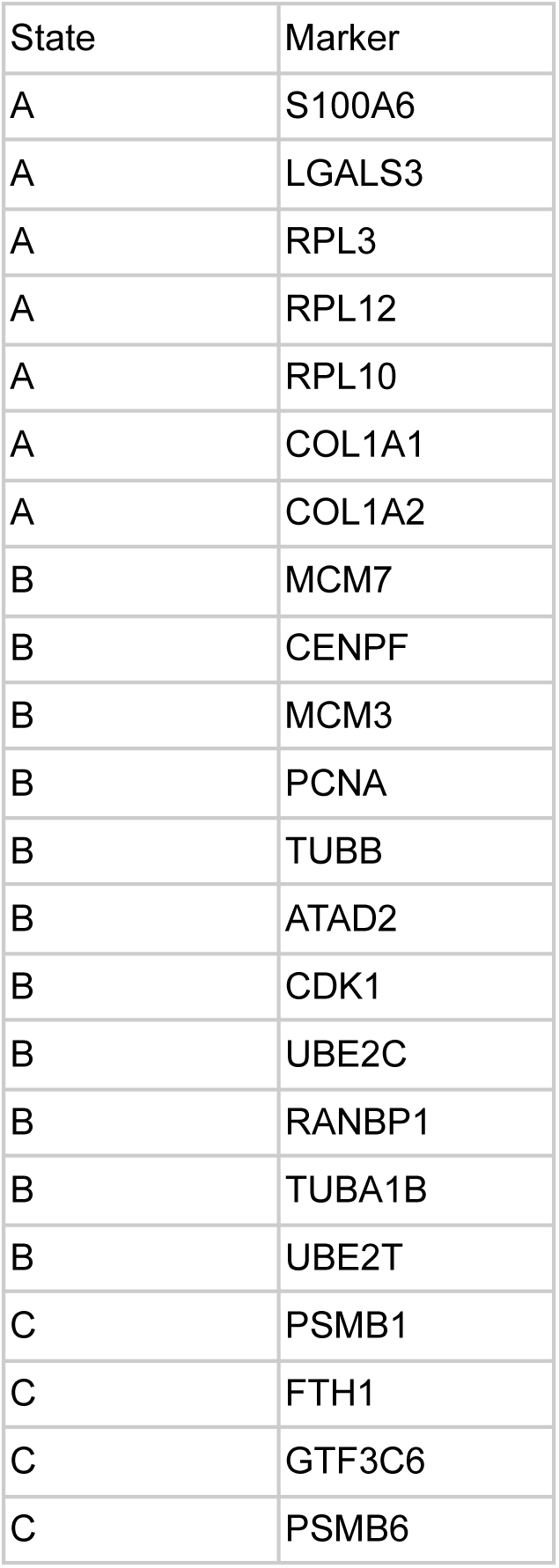

We also used GSEA to identify pathways enriched in each state. This analysis helped identify the enrichment of TGF-ß, PI3K/AKT/mTOR and VEGFA pathways in State “A” **(Figure 2E)**. The genes and pathways enriched in State “A” have previously been associated with metastatic and recurrent OS tumors.^11^ We also found that the State “A” had higher rates of cells within the G1 cell-cycle phase **(Figure S2A)**. State “B”, on the other hand, enriched for the CSR and GCNP pathways while State “C” enriched for the ESC pathway **(Figure 2E)**. We also confirmed enrichment of these State gene modules within clusters from a primary OS dataset **(Figure S1C)**.^11^ This analysis supports the existence of transcriptionally similar subpopulations across the samples and suggests the likelihood that there is significant transcriptional heterogeneity within tumors.

### Lineage tracing identifies state with increased sensitivity to therapy in OS PDX-derived cell lines

Having defined the transcriptional similarity within subpopulations present within the OS PDX-derived cell lines, we sought to determine whether the three identified states had variable sensitivity to specific drugs. To enable the identification of sensitive clones after performing scRNA-seq at baseline, we used a barcode library containing barcodes that, upon transduction, are both integrated into the genomic DNA and expressed as part of the 3’ UTR of BFP.^20^ Detection of the same barcodes in both genomic DNA and scRNAseq libraries allows for the analysis of barcode frequencies after different therapeutic conditions using the gDNA and simultaneously enables association of barcoded cells to specific transcriptomic phenotypes. Using this approach, we explored the variable sensitivity of cell states within the OS PDX-derived cell lines at a single-cell resolution.

Transduced cells were sorted followed by plating at low density to downsize the total number of recovered barcodes to approximately 1,000 **(Figure 3A)**. Cells were then treated with the IC50 of one of three drugs i) Cisplatin ii) CDK 4/6 inhibitor or iii) ATR inhibitor **(Figure 3A and S3B)**. Cisplatin was used because it is part of the chemotherapy standard of care for OS.^14^ Both CDK-4/6 and ATR inhibitors have been previously tested in OS, but found to have variable efficacy.^21,22^ Thus we hypothesized they would be useful agents to evaluate the hypothesis that modulation of cell state could increase sensitivity to these agents.

**Figure 3:**
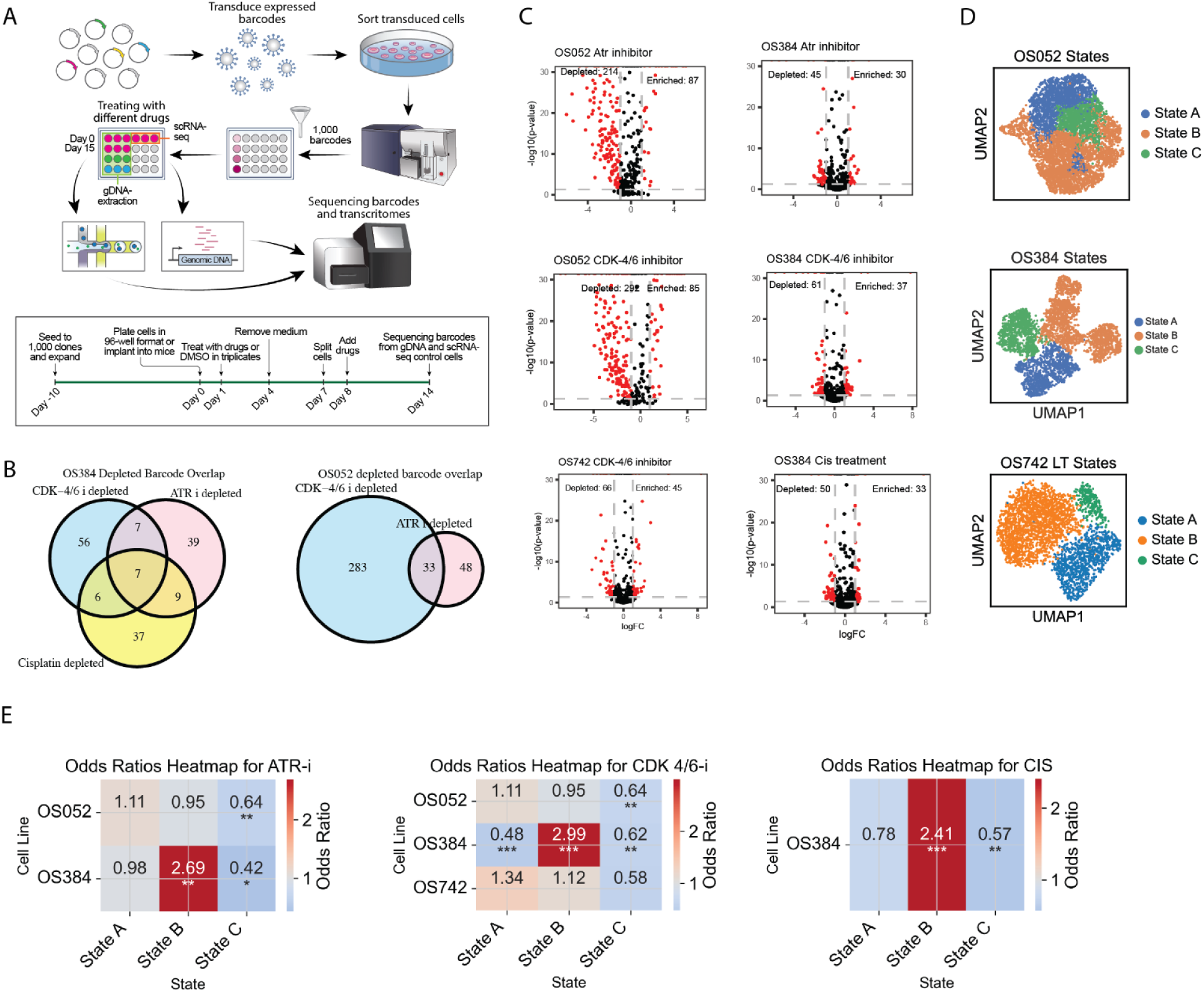
Identifying the sensitive state using lineage tracing. **(A)** Experimental pipeline for the lineage tracing approach to identify the sensitive state in the OS PDX-derived cell lines. Here, we transduced cells with lentiviral particles containing the expressed barcodes and sorted the transduced cells using blue fluorescent protein (BFP). The cells were then bottlenecked to 1,000 cells and expanded prior to being treated with different drugs. Every treatment condition in the experiment was performed on n=3 technical replicates. After treatment with a 15-day drug regimen, the amplified barcodes and single-cell libraries were sequenced. **(B)** The venn diagrams illustrate the overlap of depleted barcodes in OS384 treated with CDK-4/6 inhibitor (CDK-4/6 i), ATR inhibitor (ATR i), and Cisplatin and the overlap of depleted barcodes in OS052 with CDK 4/6 and ATR inhibitors. The overlap between each pair of depleted barcode sets is shown. Fisher’s Exact Test was used to evaluate the significance of the barcode overlaps with pairwise comparisons and adjusted p-values with Benjamini-Hochberg (CDK 4-6 i vs. ATR i p=0.000431, ATR i vs. Cis p=1.16e-06 and CDK 4/6 i vs. Cis p = 0.000662). A hypergeometric test was also performed to determine the significance of the 3-way overlap (p=0.25). **(C)** Enriched and depleted lineage tracing barcodes in OS052, OS384 and OS742. The enriched and depleted barcodes for the ATR inhibitor, a CDK 4/6 inhibitor and Cisplatin are highlighted in red and were defined as those that had p-values less than 0.05 and 𝑙𝑜𝑔_2_ fold-change less than 1 (for depleted barcodes) or more than 1 (for enriched barcodes) **(D)** Here, we are showing the scRNA-seq clustered embeddings for the untreated OS052, OS384 and OS742 cell lines after being transduced with the lineage tracing barcode library. The cells that were sequenced for this experiment included cells from n=3 technical replicates for untreated controls. **(E)** The likelihood of lineage tracing barcodes being depleted based on the State they belonged to. The heatmaps of the odds ratios are based on binary logistic regression.

After treatment with one of these three drugs, we harvested the genomic DNA from the treated and control replicate sets and amplified the LT barcodes **(Figure 3A; see Methods)**. We could then ensure that there were adequate counts for each replicate **(Figure S4A)** and compute correlations of each replicate to assess experimental precision **(Figure S4B and S1A)**. After plotting the count distributions, we found that certain barcodes contained far higher counts, suggestive of increased replicative capacity compared to other clones **(Figure S4C)**. We then plotted the ranked barcodes and filtered the dataset based on average log for the time 0 technical replicates **(Figure S4D)**. This analysis ensured that we were using high-quality counts within our analysis and provided a whitelist with which to filter the datasets from other time points and treatment conditions.

We then identified enriched and depleted barcodes by taking the log-fold-change of the counts per million (cpm) values between the treated and control samples. For example, OS052 had higher rates of depleted and enriched barcodes with more than 200 depleted and more than 80 enriched barcodes for both the ATR-inhibitor treatment and CDK-4/6 inhibitor **(Figure 3C)**. In OS384, on the other hand, we identified 61 depleted and 37 enriched barcodes for the CDK-4/6 treatment, 50 depleted and 33 enriched barcodes for the Cisplatin treatment and 45 depleted and 30 enriched for ATR treatment, suggesting variable clonal sensitivity to this therapy (**Figure 3C)**. Finally, OS742 had 66 depleted and 45 enriched barcodes for CDK-4/6 treatment **(Figure 3C)**. After comparing the depleted barcode overlap for different treatment combinations in OS384, we found that a significant proportion of barcodes were commonly depleted in different treatments **(Figure 3B)**. These findings support the hypothesis that the progeny of individual cells have differential sensitivity to therapy. Identification of the depleted and enriched barcodes also helped inform sensitive states within the cell lines after performing scRNA-sequencing on another control replicate set.

After performing scRNA-sequencing on the control replicate set **(Figure S5A)**, targeted PCR-enrichment and sequencing of the barcode-carrying BFP transcripts allowed for mapping of the LT barcodes to the associated 10X cell-barcodes. This allowed us to identify which cluster each of the enriched and depleted LT barcodes belonged to, enabling the comparison of the proportions of depleted barcodes within all cells for each state and the proportion of depleted barcodes within the rest of the cells. After designating clusters with the correct State assignment **(Figure 3D)**, we could infer if cells within a given state demonstrated higher rates of sensitization to each drug. For all samples, we found that State C had a lower likelihood of containing depleted barcodes suggesting a State with elevated resistance **(Figure 3E)**. We also found that State B in OS384 contained a higher likelihood of containing depleted barcodes (OS384: State B odds ratio of 2.99 for CDK4-6 i treatment, 2.69 for ATR treatment and 2.41 for Cisplatin treatment) **(Figure 3E)**. These findings suggest that State “B” transcriptomic profile is associated with the sensitive phenotype in OS384 and OS742 and helped identify States “A” and “C” as the states to target for reprogramming to improve therapy sensitivity. The elevated resistant phenotype in State “A” may be due to enriched PI3K/AKT/mTOR and TGF-ß pathways which are well known for mediating resistance and relapse.^23^

### Identifying State drivers

Although several pathways, such as the VEGFA and mTOR pathways, have been identified as potential targets to enhance therapeutic efficacy in OS,^6,24,25^ compensatory mechanisms can reintroduce a resistant phenotype after inhibiting these pathways.^13^ To address the limitations posed by these compensatory mechanisms, we focused on modulating cell states rather than targeting individual pathways. In doing so, we aimed to increase the proportion of cells in the sensitive state by inhibiting transcription of suspected drivers for the less sensitive states **(Figure 4A)**.

**Figure 4:**
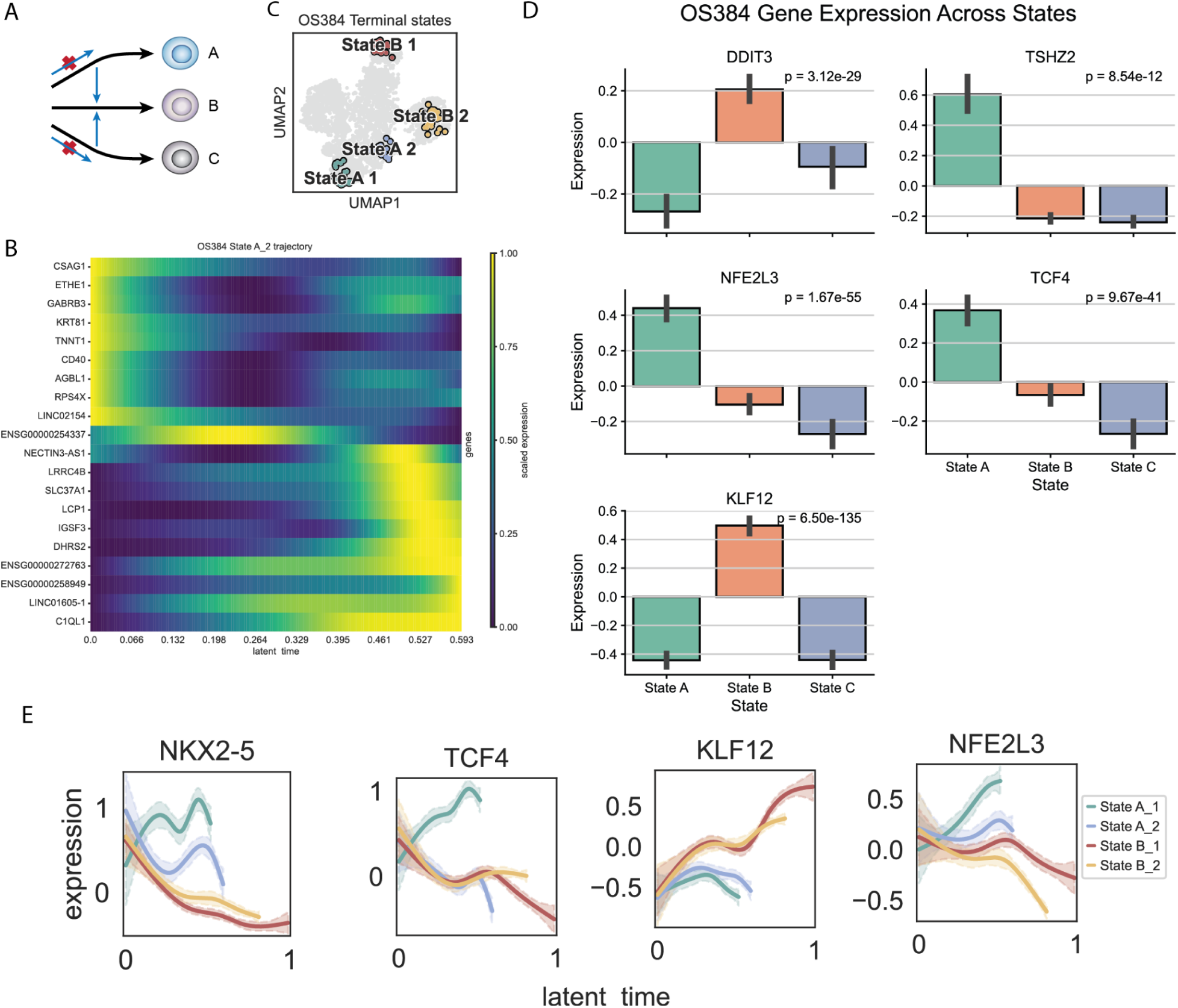
Identifying OS PDX-derived cell lineage drivers using CellRank and a biologically informed Variational Autoencoder (VAE) **(A)** The goal of this analysis is to identify lineage drivers that can reprogram cells from resistant states to those associated with more sensitive phenotypes. By identifying these critical transcription factors (TFs), we aim to modulate entire cellular states to enhance sensitivity in OS PDX-derived cell lines. **(B)** The heatmap shows the expression of the top 20 driver genes associated with the “State A_2” terminal state in OS384. Cells are ordered along the x-axis based on pseudotime progression, while rows represent driver genes identified for the selected lineage. Color intensity reflects gene expression levels, with brighter colors indicating higher expression. **(C)** Here, we show the terminal macrostates of the OS384 cell line as predicted by CellRank using the “top_n” method with four states selected. The UMAP embedding visualizes these terminal states within their associated clusters. Each distinct color represents a different terminal state. **(D)** The bar plots display the mean expression levels of the identified lineage drivers across distinct transcriptional states identified in the OS384 lineage-tracing dataset. Each bar shows the mean expression for a given state with error bars indicating the standard deviation (SD). Corrected p-values are calculated using Kruskal-Wallis tests followed by Benjamini-Hochberg correction for multiple testing, with a significance threshold of 0.05. **(E)** To confirm variable expression of the CellRank identified lineage drivers, we quantified expression of the TFs through pseudotime for the four terminal states identified by CellRank. Here, we show the expression of *NFE2L3*, *TCF4* and *NKX2-5* through pseudotime in the OS384 LT transduced cell line.

Having identified State “B” as potentially a more sensitive population **(Figure 3E)**, we sought to identify TFs that could regulate the transition to cell states **(Figure 1C)**. We identified candidate regulators of these states using CellRank. CellRank utilizes the theories of state transition and trajectory inference to infer potential drivers of a specific lineage.^26^ Recently, various tools have been released to model state transitions and trajectories with pseudotime, a unit-less time measurement.^27^ For example, after modeling gene expression through pseudotime for the State “A” trajectory, we found that *CSAG1* and *CD40* were expressed early on within the trajectory while *IGSF3* was expressed later on within the trajectory **(Figure 4B)**. Genes that are enriched early on within one trajectory relative to others may serve as drivers for the respective trajectories.

Having identified terminal states using CellRank **(Figure 4C)**, we could then identify candidate drivers for the terminal states by correlating expression of genes with fate probabilities for a given terminal state. Using this approach, we identified 10 candidate transcription factor candidate drivers, *DDIT3, TSHZ2, STAT1, NR0B1, NFE2L3, TCF4, NFKB2, IRF1, ZIC2* and *KLF12* **(Figure 4D, 4E, S6A and S6B)**.

To complement this analysis, we used CiberATAC to infer TFs suspected of regulating State “A”. CiberATAC is designed to capture regulatory interactions by integrating biological knowledge into its neural network architecture, representing regulatory interactions between TFs and their regulated genes with edges in the first two layers of an encoder. Using this approach we identified transcription factors *JUNB, DDIT3* and *FOS* as potential drivers for State “C” in multiple samples (**Figure S6C**).

These transcription factors are associated with poor prognosis in other cancers, suggesting that they could also contribute to the resistant phenotype observed in OS State “C”.^28,29^ In total, we identified 12 suspected transcription factor lineage regulators to further investigate **(Table 2)**. We then wanted to determine whether targeting these TFs could reprogram cells toward the more sensitive phenotype.

### Reprogramming cells to sensitive state and testing efficacy of top targets

The 12 TFs identified as suspected drivers were knocked down using the Perturb-seq pipeline which enables the combination of pooled screens with scRNA-seq to simultaneously capture the guide-RNA and associated transcriptome.^30^ After transducing OS384, OS742 and OS833 with the dCas9-KRAB m-Cherry and the dual-guide RNA (gRNA) constructs,^31,32^ we sorted the successfully transduced cells for both mCherry and BFP expression **(Figure S7A)**. Sorted cells were cultured for 72 hours and used to generate scRNA-seq libraries with the 10X platform **(Figure 5A)**.

**Figure 5:**
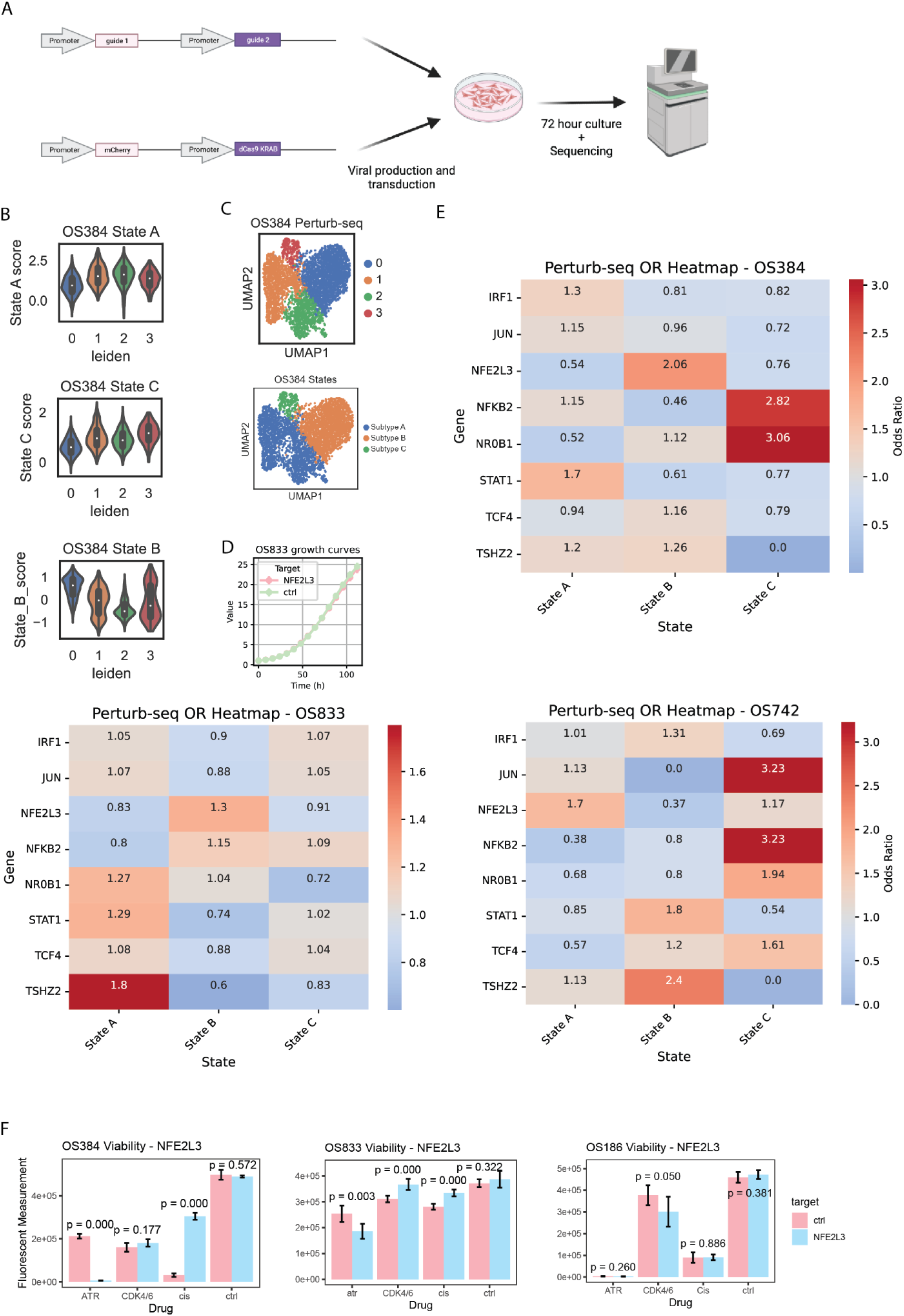
Using Perturb-seq to test reprogramming capabilities and testing efficacy improvements after perturbations. **(A)** Experimental pipeline for the Perturb-seq experiment. After generating three OS-PDX derived cell lines co-expressing mCherry and dCas9-KRAB (CRISPRi), we transduced the OS PDX-derived cell lines with lentiviral particles containing constructs with dual-guide RNAs for the same target in addition to BFP. The cells were then sorted for double positive mCherry and BFP and processed with the 10X 3’ v3.1 kit **(B)** State gene module enrichment in the different OS384 clusters. **(C)** Embedding for the OS384 perturb-seq sample. These cells were transduced with the perturb-seq library. On the left, we are showing enrichment of the State B module in the different clusters identified using the Leiden cluster algorithm. **(D)** We compared the growth rates of the NFE2L3 knockdown cell line and control cell line by plating the cells with n=6 technical replicates in a 96 well format and quantifying confluency with the incucyte. The control cell line is highlighted in green while the NFE2L3 knockdown cell line is highlighted in red. The error bars represent standard deviations for the technical replicates. **(E)** To determine whether the knockdown of the TFs in the perturb-seq library increased the likelihood of cells being in a certain state using binary logistic regression models within each cell line. Here we are showing the log odds ratio for the different TFs. **(F)** To test sensitivity differences to an ATR inhibitor after knockdown of NFE2L3, we treated the control and NFE2L3-KD cells with the 15 day drug regimen and quantified viability with the CellTiter Glo assay. Error bars represent standard deviations for the n=3 technical replicates and the p values were computed using a Student’s t-test.

After analyzing this scRNA-seq dataset and confirming proper knockdown of TFs **(Figure S8B)**, we identified which perturbations increased the proportion of cells in each state. To do so, we filtered the datasets based on quality control parameters **(Figure S7B; see Methods)** and quantified changes in state proportions for each perturbation. After confirming that certain clusters enriched for the State modules, we classified each cluster to a certain state by computing the correlation between average cluster expression values and those of the previously NMF defined states for OS152, OS384 and OS742 **(Figure 5B and 5C)**.

To evaluate reprogramming capabilities of the candidate TF drivers, we used binary logistic regression models to predict the likelihood of cells being in a certain State based on the TF that was targeted with the cell line variable used as a covariate. We found that *NFE2L3* knock-down (KD) resulted in higher likelihoods of cells being in State B for OS384 and OS833. Interestingly, the *NFE2L3* KD also increased the likelihood of OS742 cells being in State A, which was previously identified as the sensitive state for OS742. *TSHZ2* KD also reduced the likelihood of cells being in State “C” **(Figure 5E)**. *NFE2L3* was found to promote radiotherapy resistance in esophageal squamous cell carcinoma and found to regulate resistance via the *IL-6* pathway.^33–35^

After confirming *NFE2L3* knockdown **(Figure S8E)**, we tested treatment sensitivity after knocking down these targets with the dCas9/CRISPRi system. *NFE2L3* KD and control cells were treated with Cisplatin, an ATR inhibitor and a CDK-4/6 inhibitor with the same 15-day drug regimen used in the LT experiment and viability was quantified using the CellTiterGlo assay. *NFE2L3* KD improved the efficacy of the ATR inhibitor in both OS384 and OS833 and improved the efficacy of the CDK-4/6 inhibitor in OS186. Surprisingly, KD of NFE2L3 seemed to decrease the sensitivity to cisplatin (p = 0.003, t-test, **Figure 5F)**. We found no differences in the growth rates between the *NFE2L3* KD and control cells **(Figure 5D)**. To explore the potential mechanisms for this phenotype, we investigated the downstream modulated genes after *NFE2L3* knockdown. Knockdown of *NFE2L3* resulted in downregulated expression of *TGFß1* and *KIF1A* **(Figure S8C)**, potential mechanisms for sensitization within these samples. TGF-ß, for example, has been identified as a compensatory mechanism for BRAF, EGFR and HER2 inhibition.^36,37^ Target genes for NFE2L3 were also downregulated in NFE2L3 KD cells suggesting that the NFE2L3 regulome was effectively modulated **(Figure S8D)**.

## Discussion

Heterogeneity and compensatory mechanisms remain two of the most difficult challenges for successful treatment of various cancers. Osteosarcoma (OS), an aggressive pediatric cancer characterized by early metastasis and significant treatment resistance, demonstrates these same challenges.^38^ The survival rate for patients with metastatic OS remains below 30%.^39^ Resistance to targeted therapy in OS and many other cancers is likely due to extensive intratumor heterogeneity and the activation of compensatory survival pathways.^11,40,41^ These hurdles underscore the importance of studying resistance from a systems perspective by assaying entire transcriptomic states.^5,20^ Several pathways, including mTOR, TGF-ß, and VEGFA, have been previously implicated in OS resistance, many of which are particularly enriched in metastatic cells, exacerbating the treatment challenges and contributing to the poor survival rates observed in patients with metastatic disease.^43^ In fact, pathways such as TGF-ß has been recognized in so many cancers as promoting angiogenesis and suppressing anti-tumor immunity, that various groups have developed ways to inhibit TGF-ß.^44^ Our study represents a shift in this paradigm by identifying and targeting cellular states rather than individual pathways, thereby offering a more systematic approach to combating resistance.

We identified three states within the OS PDX-derived cells lines and the sensitive state using a LT approach. We identified State C as the more resistant and State B as the more sensitive states. After identifying and perturbing suspected drivers for the specific states using the perturb-seq pipeline, we found that *NFE2L3* KD resulted in higher likelihoods of cells being in State B. *NFE2L3* has previously been identified as a potential target in Bladder Cancer and hepatocellular carcinoma which promotes cell proliferation.^45,46^ We also found that *NFE2L3* KD was able to sensitize the OS PDX-derived cell lines to an ATR inhibitor, but not to cisplatin or a CDK-4/6 inhibitor. In fact, the *NFE2L3* KD seemed to make the cells more resistant to cisplatin.

Using this strategy, we show how intra-sample heterogeneity can be analyzed to identify targets capable of shifting an increased proportion of cancer cells to a sensitive state. We were able to modulate multiple pathways at once and reduce the likelihood of compensatory pathways reintroducing resistance — a common problem in Osteosarcoma and other cancers. This approach could be used to improve the efficacy of various emerging therapies including immunotherapies.^47^

While individual perturbations like knocking down *NFE2L3* has demonstrated potential in shifting cells towards a sensitive state in our study, it is increasingly evident that combinatorial perturbations may yield more robust outcomes for reprogramming purposes.^48^ The underlying rationale is that by targeting multiple transcription factors, it is possible to interfere with the compensatory mechanisms that often undermine the efficacy of single-agent therapies.

However, the exploration of combinatorial perturbations introduces significant challenges, due to increasing costs with the vast number of potential combinations. To address this, recent tools offer promising avenues for predicting the outcomes of multi-target perturbations without the need for exhaustive empirical testing. Tools such as CPA, CellOracle, and GEARS have shown efficacy in simulating transcriptomic changes post-perturbation, enabling the identification of potentially efficacious combinations.^49,50^

By integrating these tools into our research pipeline, we can systematically explore the combinatorial landscape, effectively narrowing down the options to those most likely to induce a reprogrammed, sensitive state in osteosarcoma cells. Moreover, novel constructs capable of delivering multiple gRNAs in a single intervention have recently been introduced and can further enhance the practical applicability of these combinatorial approaches,^51^ paving the way for the modulation of even more pathways simultaneously and perhaps further improving therapeutic efficacy in various cancers.

## Limitations of the Study

A major limitation of this study may be the inherent plasticity of the cells, allowing them to transition to a more resistant state after a therapeutic inhibitor degrades. This limitation could perhaps be addressed by the use of multiple doses or the gradual release of the therapeutic agent for a more consistent inhibition of the target. Efficacy improvements may also be mitigated by the gradual induction of compensatory mechanisms, necessitating longitudinal studies to identify resistance mechanisms that arise after a given amount of time.

Although this approach successfully reprogrammed a large proportion of the cells, we may be able to reprogram a larger proportion by targeting TFs that might reprogram cells out of terminal states as well. Despite these limitations, the approach taken in this study may provide a foundation for sensitizing tumor cells to existing or novel therapies.

### Reagents table

| Reagent or Resource | Company | ISBN/Cat. number |
| --- | --- | --- |
| DMEM medium | Gibco |  |
| Agilent tapestation reagents | Avantor | 76645-270 |
| TransIT-293 Transfection Reagent | Mirus Bio | MIR 2705 |
| DNeasy Blood and Tissue Kit | Qiagen | 57205964 |
| 10X kit 3' V3.1 | 10X Genomics | 1000268 |
| PBS pH 7.4 (1X) | Gibco |  |
| DNA clean and concentrator | Zymo research | D4014 |
| One Shot™ Stbl3™ Chemically Competent E. coli | Thermo Fisher Scientific | C737303 |
| White walled, clear bottom 96-well plate | Greiner | 655098 |
| Cell Titer Glo | Promega | G9241 |
| Elimusertib (BAY-1895344) hydrochloride (ATR inhibitor) | Selleckchem | S8666 |
| Q5 Hot Start hifi master 2X master mix | New England Biolabs | M0494S |
| Zymoclean Gel DNA Recovery KIT | Zymo research | D4007 |
| Opti-MEM I (1X) | gibco | 31985-070 |

## Methods

### Cell Cultures

The OS PDX-derived cell lines OS052, OS152, OS384, OS742 and OS833 were used for the experiments in this study and were originally derived from PDX models.^52^ The cell lines were generated by implanting patient Osteosarcoma (OS) tumor cells in mice and then culturing the cells from the resected mouse tumors.^17^ The cell lines were cultured for less than 13 passages in DMEM +Glutamine (Gibco) with 5% FBS and Penicillin-Streptomycin. HEK293T cells were also cultured in DMEM +Glutamine (Gibco) with 5% FBS and Penicillin-Streptomycin. All cells were kept in a 37 °C and 5 % humidified CO_2_ incubator.

### Single-cell RNA sequencing, quality control and read-alignment

10X scRNA-seq libraries were prepared with the v3.1 kit. Prior to GEM preparation, 10,000 cells were prepared in cold PBS and loaded onto a Next GEM chip before being loaded onto the 10X Chromium X. After reverse transcription, the cDNA libraries were amplified for 11 cycles, analyzed on the Agilent Tapestation and quantified on the Qubit before being sequenced on the Novaseq machine. The single-cell RNA sequencing FASTQ files were then processed using Cell Ranger version-7 with default settings. The “feature-reference” argument was used for the Perturb-seq data to assign LT barcodes and gRNAs respectively to individual cells. Cellsorted-possorted bam files were then generated with Samtools to be used by velocyto to generate loom files.^53^

After creating an anndata object from the loom files with Scanpy, the data was filtered based on the number of genes, total counts and mitochondrial percentage before being regressed on mitochondrial percentages and ccdifference scores. The ccdifference scores were computed by taking the difference between the G2M and S phase scores which were generated by quantifying enrichment of G2 and S gene lists with the Scanpy ‘score_cell_cycle_genes’ command.^54^ The same preprocessing pipeline was applied to the Zhou dataset.^11^

Prior to umap generation, dimensionality reduction and clustering were performed with PCA and the leiden algorithm, respectively. Prior to assigning clusters to states, we aggregated the clustered datasets from the different samples, We then computed the average expression for each cluster and gene and normalized that data using z-score normalization. We could then aggregate the different clusters using non-negative matrix factorization (NMF), assigning clusters to one of the 3 groups based on the dominant component weights. After identifying markers for all states within all samples using the rank_genes_groups Scanpy command, we took the overlap of the markers from transcriptionally similar states across samples. Those markers that were present in two or more of the samples were then used to define gene State modules “A”, “B” and “C”.

A chi-squared test was used to determine the significance of association between cell cycle phase and state. Gene set enrichment analysis was then conducted with the Scanpy ‘score_genes’ command. The C6 oncogenic gene set collections from MSigDB were used for the gene set enrichment analysis.

### RNA velocity and trajectory inference

Using scVelo, RNA velocity analysis was performed to identify trajectories within the OS PDX-derived cell lines^55^. scVelo, a recent RNA velocity tool, models transcription, splicing and degradation rates using the dynamical model.^56^ RNA velocity models transcript kinetics to ultimately infer a cell’s state transition pattern, allowing for the inference of sources and terminal states within an embedding.

### Transducing cells with lineage tracing construct

The LT construct used in this study was derived from the original perturb-seq vector which contained a barcode diversity of ∼100,000^57^ which contained 18-mer barcodes expressed on the 3’ UTR of BFP (pBA571). We generated lentiviral particles carrying the LT barcode library by transfecting HEK293T cells with pBA571. We subsequently transduced 3 × 10⁶ cells of each cell line with these lentiviral particles. During this transduction process, the cells were cultured in DMEM + Glutamine with 5% BGS and a penicillin-streptomycin supplement. After the cells were infected at a low UMI (∼7%) the BFP+ cells were sorted using the FACS Aria. The cells were then bottlenecked and expanded ∼30 fold before being distributed into 18 wells to be treated with different drugs or with a DMSO control.

### IC50 experiments

IC50 experiments for Cisplatin, an ATR 1895344 inhibitor and CDK 4/6 inhibitor were performed for all cell lines in a 96 well format in triplicates. Starting concentrations were 5 μM for Cisplatin, 0.5 μM for the ATR inhibitor, and 2 μM for the CDK4/6 inhibitor. Eight serial 1:3 dilutions were prepared for each drug in culture medium, with the highest DMSO concentration serving as the vehicle control.

After quantifying viability with CellTiterGlo (Promega, G9241) and the BioTek Synergy neo2 plate reader, the IC₅₀ values were determined by fitting the data to a four-parameter logistic (4PL) model using the drm function from the’drc’ package in R.

### FACS sorting

The cells used for the LT and Perturb-seq experiments were first spun down at 300g for 5 min and resuspended in cold PBS with 10% FBS. After the cells were passed through a 70*u*M filter and placed on ice, they were sorted for BFP and mCherry double positive signal using the BD FACS Aria and sorted into 50% DMEM medium and 50% FBS.

### Treating the lineage tracing transduced cells and sequencing the barcodes from genomic DNA

The 15-day drug regimen included 3 days of treatment, a 3-day drug holiday and 7 more days of treatment. The 3-day drug holiday was included to allow resistant clones to proliferate.

Genomic DNA (gDNA) from the LT experiment was extracted using the DNeasy Blood and Tissue Kit (57205964, Qiagen). The barcodes were then amplified from the gDNA using Primer 1 and 2 for 25 cycles with an annealing temp of 71C with the KAPA HIFI master mix. The first PCR isolated the barcode from the gDNA and added on the Truseq read 1 and read 2 primer binding sequences. A second PCR was then performed to append the unique dual indices and P5 and P7 adapters to allow for Illumina sequencing. An annealing temp of 72C and 25 cycles was used for the second PCR. The barcodes were then assayed on the Agilent tape station and quantified on the Qubit 3.0 before being sequenced on the Illumina HiSeq or NovaSeq.

Primer 1: ACACTCTTTCCCTACACGACGCTCTTCCGATCTGCACAGTCGAGGCTGAT

Primer 2: GTGACTGGAGTTCAGACGTGTGCTCTTCCGATCTCAAACTGGGGCACAAGC

### Lineage tracing barcode analysis

The whitelist for the LT barcodes was created from the Day-0 control sample by plotting the ranked counts and designating a threshold close to the elbow to filter the dataset to 988 barcodes for OS384, 961 barcodes for OS742 and 1,216 barcodes for OS052. Additional barcode counts were added if the barcodes were within a hamming distance of one (hamming distance of one represents a difference of one base pair) from any barcode in the whitelist. We then filtered the barcode list in all the other conditions based on the whitelist and added values from non-whitelist barcodes that were within a hamming distance of one. To compare barcode representation in different conditions, we computed the log-fold change of the cpm scaled values between the treated samples and the day-13 control.

The p-value for this analysis was computed using Pearson’s chi-squared test with the assumption that the presence of one barcode does not influence the presence of another. The likelihood of LT barcode depletion based on the state was determined using a binary logistic regression model for each available combination of drug and cell line.

### Trajectory driver identification

CellRank was used to infer terminal states and suspected lineage drivers.^58^ The CellRank kernel employed in this study uses RNA velocity to infer directionality of the edges, representative of transition probabilities, within a markov chain. GPCCA, a CellRank estimator, was used to infer suspected drivers by correlating expression through pseudotime for each gene with absorption probabilities for a given terminal state. These inferred drivers were filtered for p-values<0.05, correlations>0.2 and based on whether they were transcription factors.

A biologically informed VAE (CiberATAC) was also used to identify suspected regulators for the state “A” profile. The nodes within the first hidden layer of the CiberATAC model are representative of transcription factors. Larger weights between the first and second layer suggest a more active transcription factor within that batch of data. We examined cell-type-specific chromatin accessibility peaks according to CiberATAC predictions and assigned each peak to the cell type with the highest observed model score. Next, we examined the over-representation of each TF within the specific peaks for each of the cell types, using a one-sided Fisher’s exact test. We reported the odds ratio and p-value for over-representation Fisher exact tests.

### Perturb-seq experiment

The dual-guide vectors used for the Perturb-seq screen in this study were assembled with a 2-step method based on the protocol detailed in the Replogle manuscript.^31^ First, ultramers containing two guide RNAs were designed for each of the 12 targets and cloned into pJR85 using Gibson cloning. These cloned constructs and pJR89 were then simultaneously digested, allowing for the tracr and promoter from pJR89, to be cloned into the intermediate construct using the Golden gate cloning method.

*5’ Gibson overhang of ultramer: tgagactataagtatcccttggagaaccaccttgttgg*

*3’ Gibson overhang of ultramer: gtttaagagctaagctggaaacagcatagcaagtttaaataa*

*Example Ultramer Sequence: tgagactataagtatcccttggagaaccaccttgttggggcaccgtgcctggcaccgcgtttcagagcgagacgtgcctgcaggatacgtctcagaaacatggggatcgggagaagcgaagagtttaagagctaagctggaaacagcatagcaagtttaaataa*

After generating viral particles for the dCas9-KRAB mCherry construct (Addgene 188779), cell lines were transduced and sorted based on mCherry expression.^32^ The sorted cells were then transduced with constructs expressing dual guide RNAs for single targets and sorted for both BFP and mCherry expression. After culturing the cells for 3-5 days, the single cell libraries were then prepared with the 10X 3’ V3.1 kit.

### Analyzing Perturb-seq dataset

Since the Perturb-seq transcript and gRNA libraries were submitted separately for sequencing, we received separate FASTQ files for each. This data was processed with cellranger with the –feature-ref and –libraries flag. We then associated the gRNA with the 10X cell barcode using the molecule_info.h5 object from the cellranger output.

We used a chi-squared analysis to test the statistical significance of cell proportions within different clusters in the context of different transcription factor knock-downs. In order to aggregate all of the samples into a single analysis, we also used a binary logistic regression model to predict the likelihood of cells being in State B given the transcription factor that was targeted for a certain cell and used the cell line as a covariate in the model.

### Testing treatment efficacy improvements for single-target knockdowns

After generating virus for NFE2L3 and NR0B1 targets, the cell lines were transduced in 10 cm dishes and FACS sorted based on BFP and mCherry into 50% DMEM and 50% FBS. The cell lines were then plated in a 96-well format in triplicates and treated with the IC50 concentrations of Cisplatin, an ATR inhibitor, a CDK 4/6 inhibitor or DMSO for the same 15-day regimen as used for the LT experiment. We use the Cell Titer Glo (CTG) (Promega, G9241) viability assay to determine any efficacy improvements with the knockdowns. After the 15-day treatment, the cells were incubated with the CTG reagent for 30 min and the fluorescence was measured using the BioTek Synergy neo2 plate reader.

Proper knockdown of *NFE2L3* for the viability studies was assessed using qPCR. The RNA was isolated using Trizol reagent followed by purification with the Zymo RNA Miniprep kit. cDNA synthesis and qPCR amplification were carried out using the Maxima reverse transcription master mix and Perfecta qPCR master mix. Fluorescence data were acquired on a Bio-Rad CFX real-time PCR instrument.

## Code Availability

https://github.com/brendamelano/Reprogramming_Osteosarcoma

## Acknowledgements

EASC was funded by the Alex’s Lemonade Foundation (Crazy 8 Initiative). HG is supported by the Arc Institute and this study was supported in part by the Arc Institute. We would also like to acknowledge Marina Sirota, Mike Keiser, Katie Pollard, Luke Gilbert and Iwei Yeh for their feedback on this project. We would also like to thank Brian Plosky and Chiara Ricci-Tam for editing and offering feedback on the manuscript.

## Supplemental Figures

**Figure S1:**
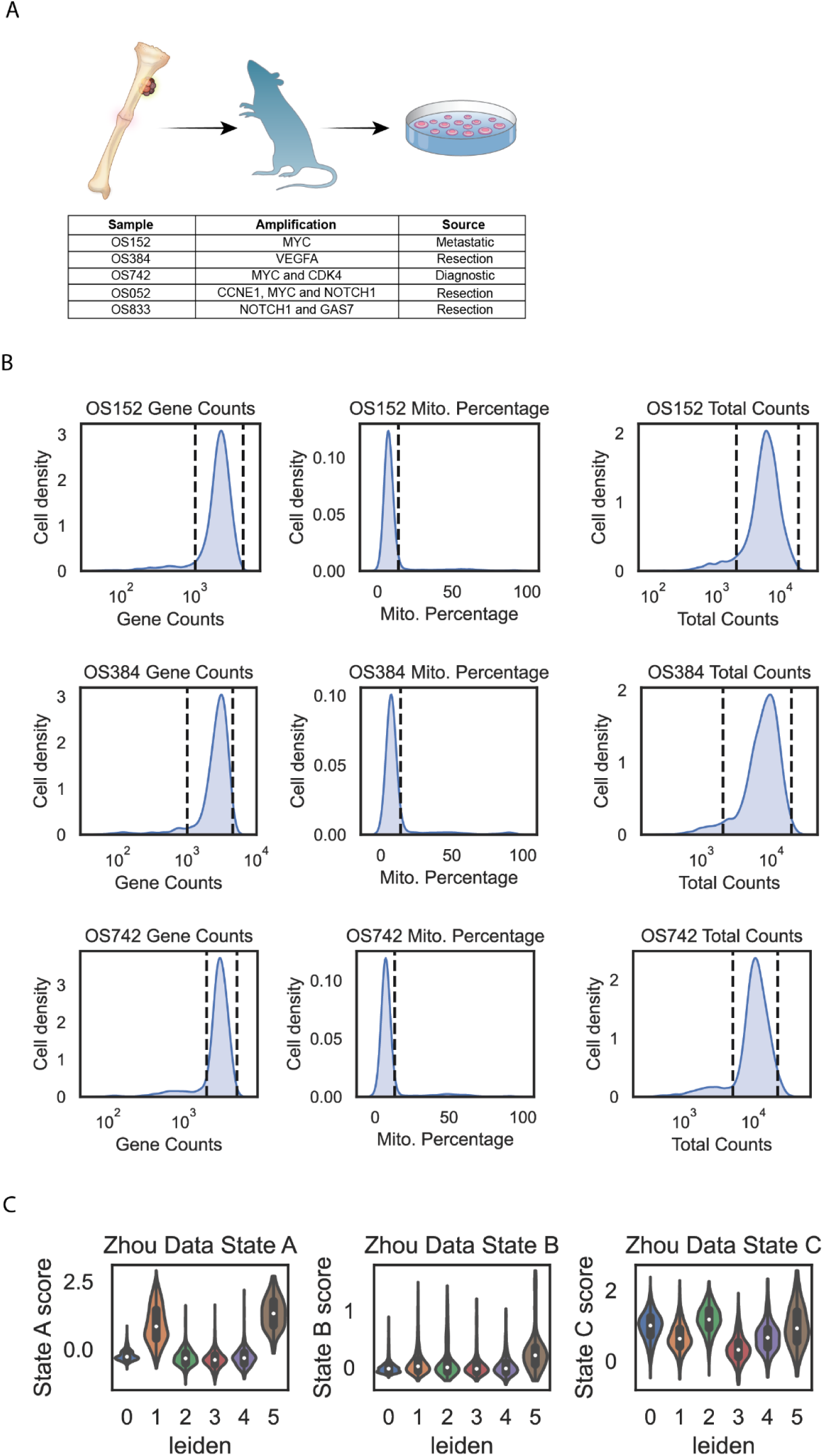
Sample descriptions and quality control analysis for scRNA-seq of OS PDX-derived cell lines. **(A)** Overview of three osteosarcoma (OS) PDX-derived cell lines—OS152, OS384, and OS742—each exhibiting amplifications in MYC, VEGFA, and CDK4. These cell lines originated from different sources and were selected for single-cell RNA-sequencing (scRNA-seq) analysis. **(B)** Quality control filtering applied to the scRNA-seq datasets, based on total read counts, number of detected genes per cell, and mitochondrial gene expression percentages. **(C)** State module enrichment for leiden clusters within a primary OS sample dataset.

**Figure S2:**
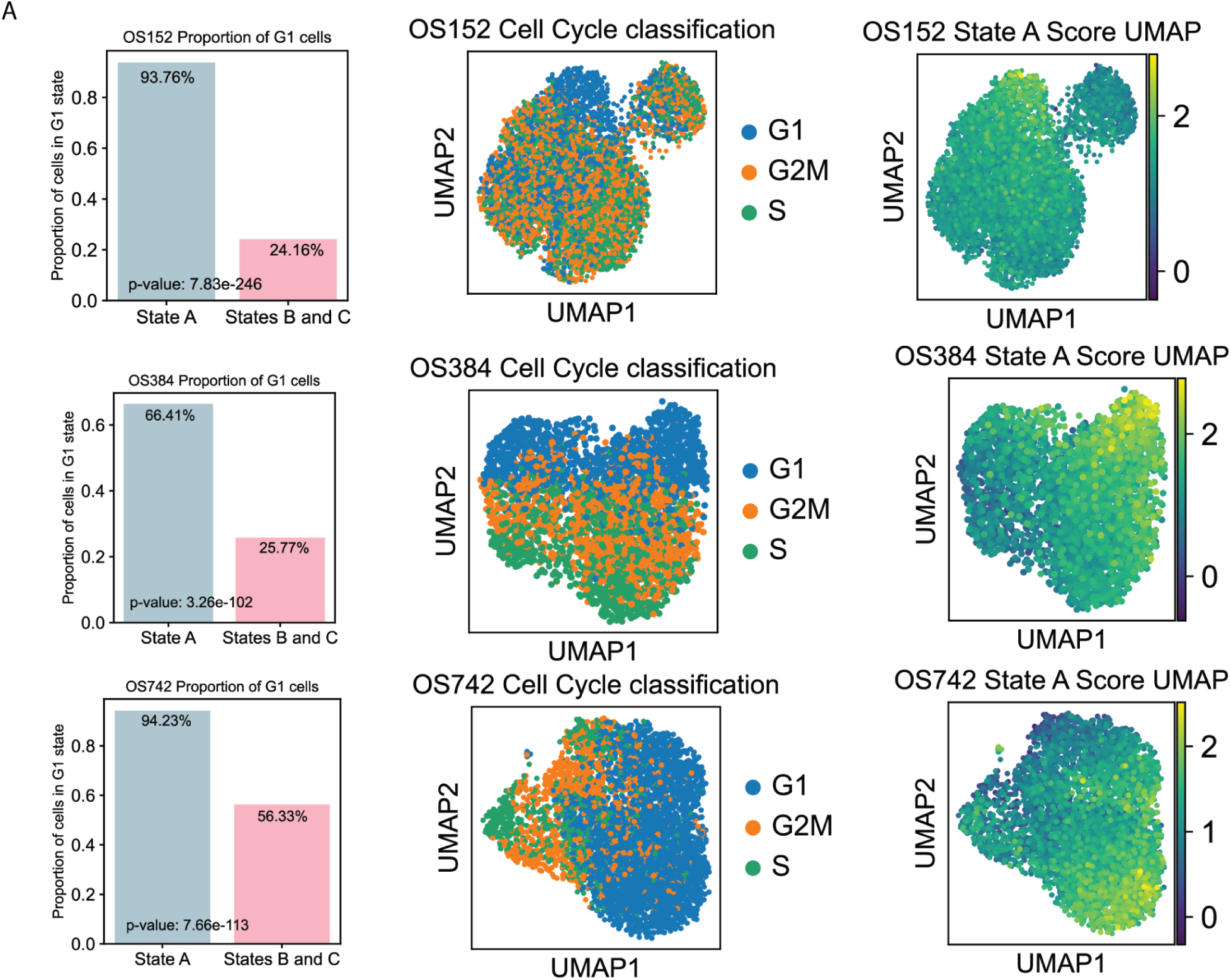
Demonstrating sample level transcriptional similarities with State gene module marker enrichment and a label transferring method. **(A)** OS152, OS384 and OS742 UMAP embeddings colored for cell cycle phase and State A module enrichment score. A chi-squared test was used to determine the significant association between the G1 phase and the State “A” population.

**Supplementary Figure 3:**
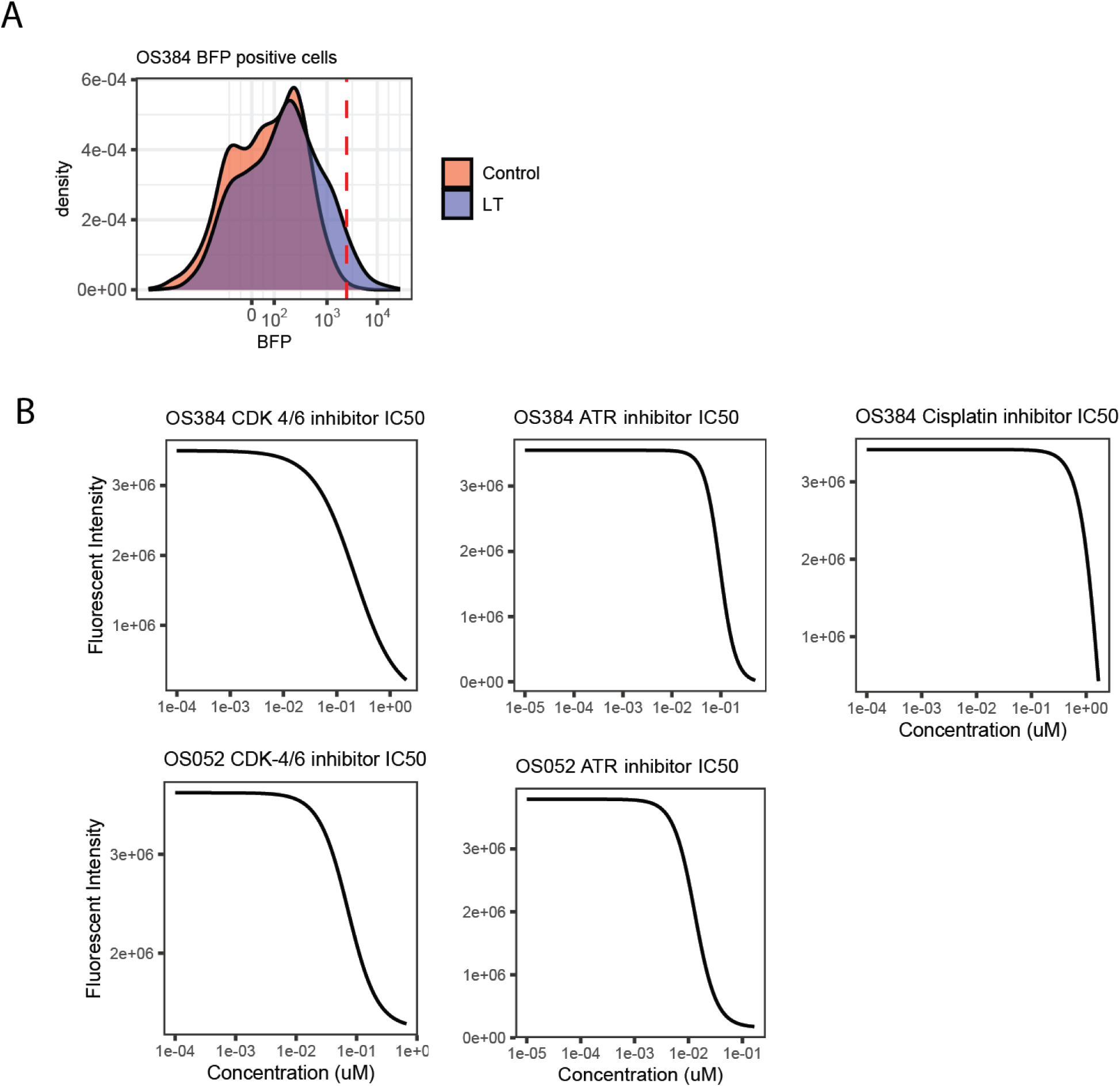
Lineage tracing sort and dose-response analysis for 15 day drug regimen used for the Lineage tracing experiment. **(A)** The overlapping histogram plots represent the BFP intensity for OS384 wild-type and lineage tracing transduced cells. Lineage tracing transduced OS384 cells are represented with the blue curve while the control cells are represented by the red curve. **(B)** The IC50 values for all drugs used on OS384 cells were determined using a dose-response assay. Cell viability was measured at varying concentrations over a 15-day period, and data were collected in triplicate. The data were fit using a four-parameter logistic regression model (LL.4), with the Hill slope, minimum, maximum, and EC50 as parameters. Fluorescent intensity was used as a proxy for cell viability. The x-axis is presented on a logarithmic scale to better visualize the concentration range.

**Supplementary Figure 4:**
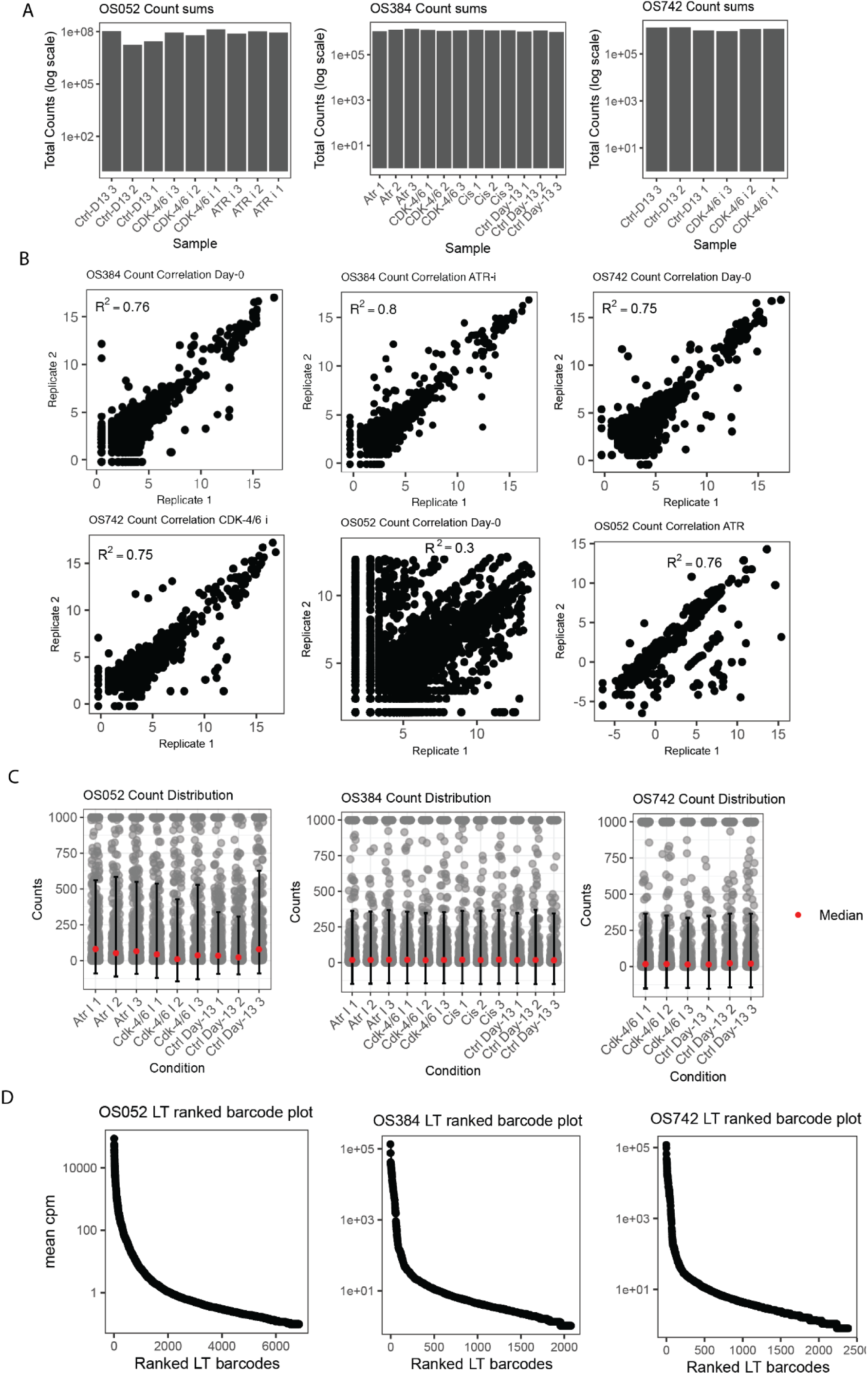
Lineage tracing sort and QC for barcodes amplified from gDNA. **(A)** Total raw counts for all replicates within OS052 and OS384 after filtering on the whitelist generated from the Day-0 samples. **(B)** Linear regression was used to compute the 𝑅^2^ for the correlation between the technical replicates for the cpm-scaled and log-transformed barcode counts. **(C)** Here, we plot the distribution of barcode counts across various conditions for OS052 and OS384, with individual data points (gray), median values (red), and error bars representing the mean ± standard deviation. Counts were capped at 1,000 to reduce the influence of extreme outliers, and x-axis labels were adjusted for clarity. Conditions include “Ctrl Day-13”, “CDK-4/6 i”, and “Atr i”. **(D)** The plots depict the ranked abundance of lineage-traced (LT) barcodes based on the mean CPM (counts per million) values for each barcode in each osteosarcomaPDX-derived cell-line. The x-axis represents the ranked LT barcodes, while the y-axis shows the mean CPM values on a logarithmic scale.

**Supplementary Figure 5:**
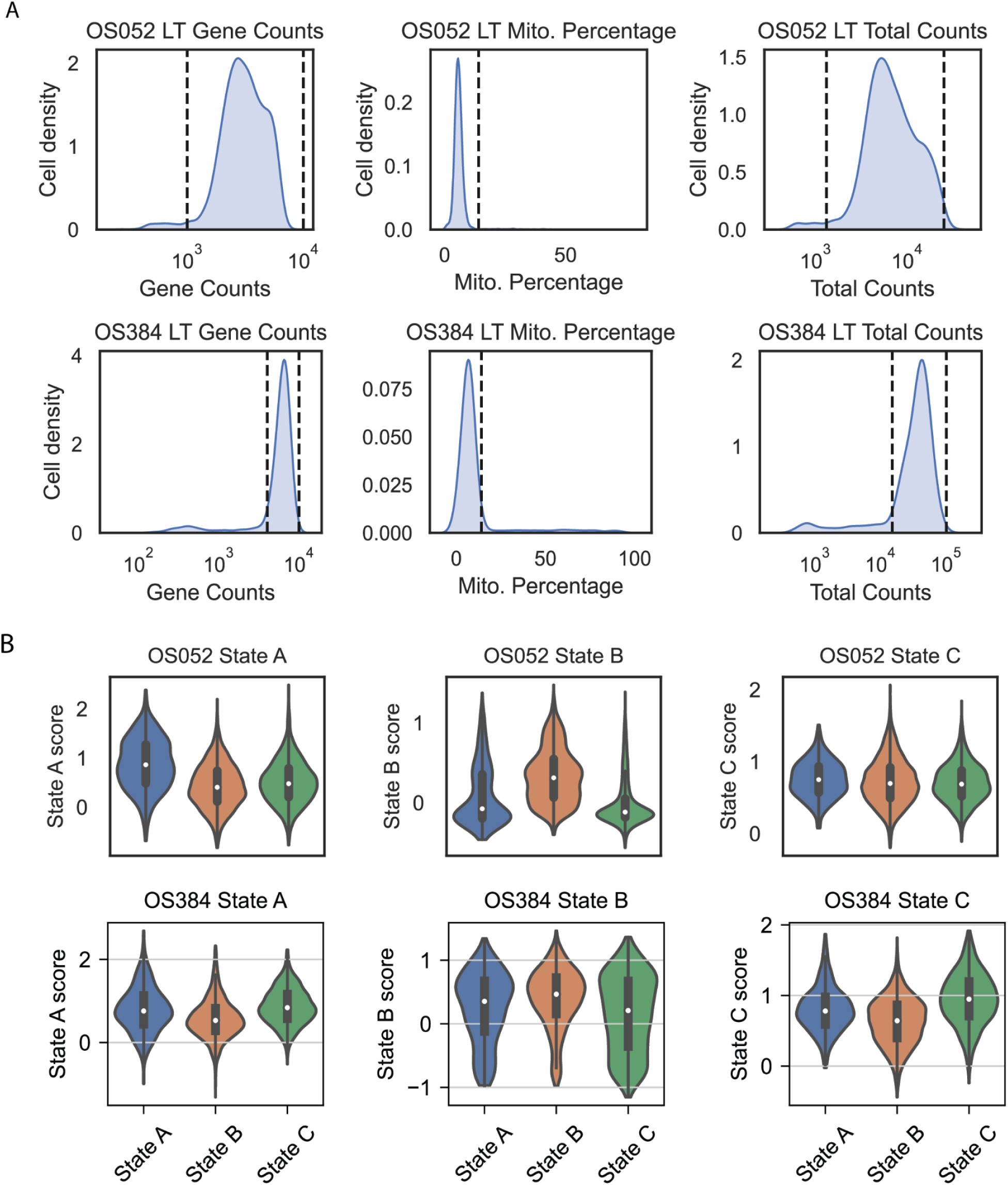
scRNAseq QC for lineage tracing experiment. **(A)** The plots show the distribution of quality control metrics for the OS052 and OS384 samples, including the number of detected genes (Gene Counts), mitochondrial gene expression percentage (Mito. Percentage), and total RNA counts per cell. Vertical dashed lines indicate threshold values used to filter out cells based on each metric. **(B)** State module enrichment in different States for the OS384 and OS052 lineage tracing transduced PDX-derived cell line.

**Supplementary Figure 6:**
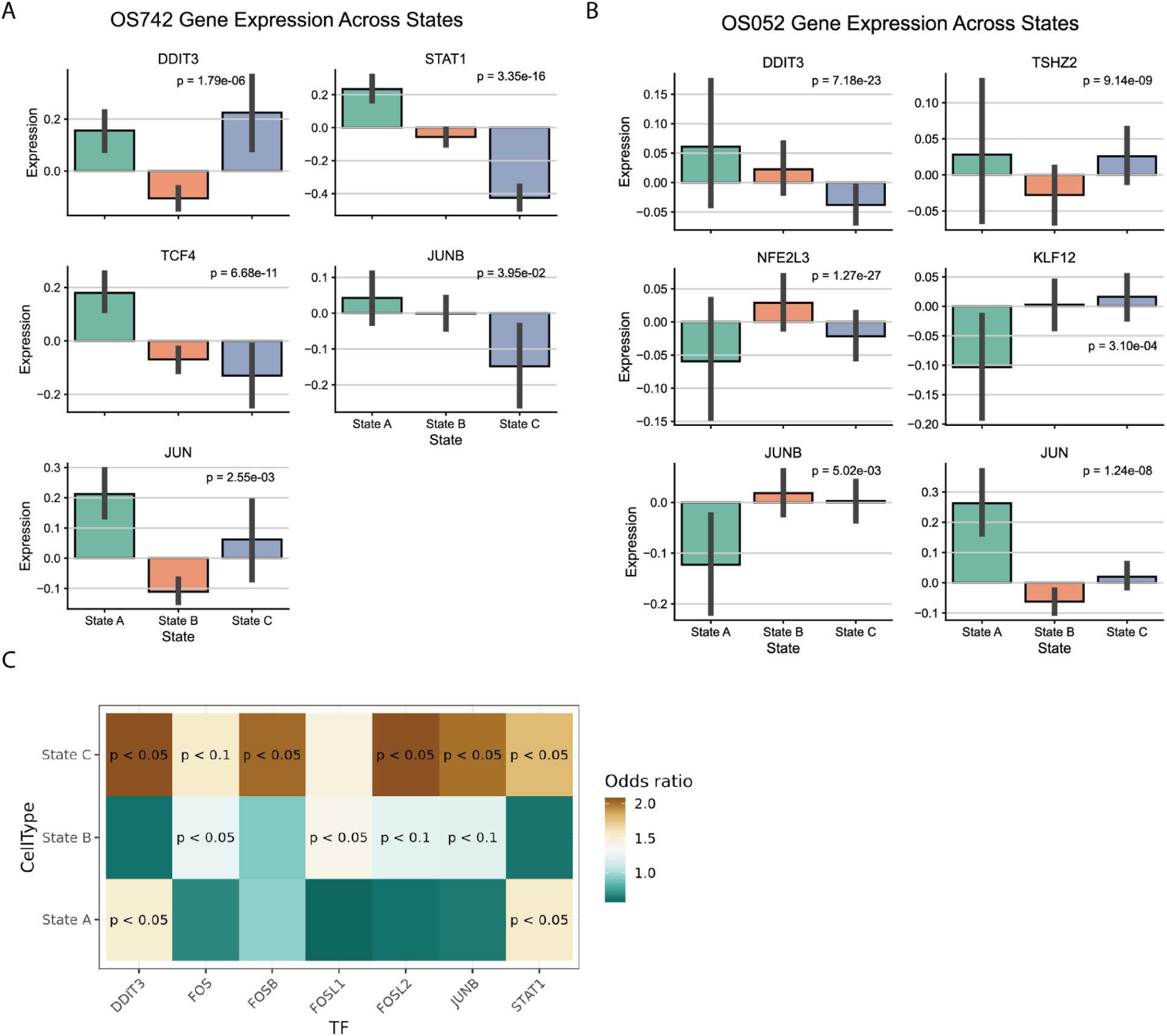
Identifying suspected drivers for States “A”. **(A)** The expression levels of selected genes were quantified and visualized across different transcriptional states in OS742 using a bar plot for each gene. Each panel corresponds to an individual gene (e.g., *DDIT3*, *STAT1*, *NR0B1*, etc.), and the x-axis represents the States “A”, “B” and “C”. Bars show the mean expression levels with standard error bars. Statistical significance between states was assessed using the Kruskal-Wallis test, and p-values were corrected for multiple testing using the Benjamini-Hochberg (BH) method. The corrected p-values are indicated in each panel. **(B)** The expression levels of selected genes were quantified and visualized across different transcriptional states in OS052 using a bar plot for each gene. Each panel corresponds to an individual gene (e.g., *DDIT3*, *STAT1*, *NR0B1*, etc.), and the x-axis represents the States “A”, “B” and “C”. Bars show the mean expression levels with standard error bars. Statistical significance between states was assessed using the Kruskal-Wallis test, and p-values were corrected for multiple testing using the Benjamini-Hochberg (BH) method. The corrected p-values are indicated in each panel. **(C)** State-specific transcription factor regulators as identified by CiberATAC, a biologically informed VAE. We are showing transcription factor odds ratio represent overrepresentation in a given state.

**Supplementary Figure 7:**
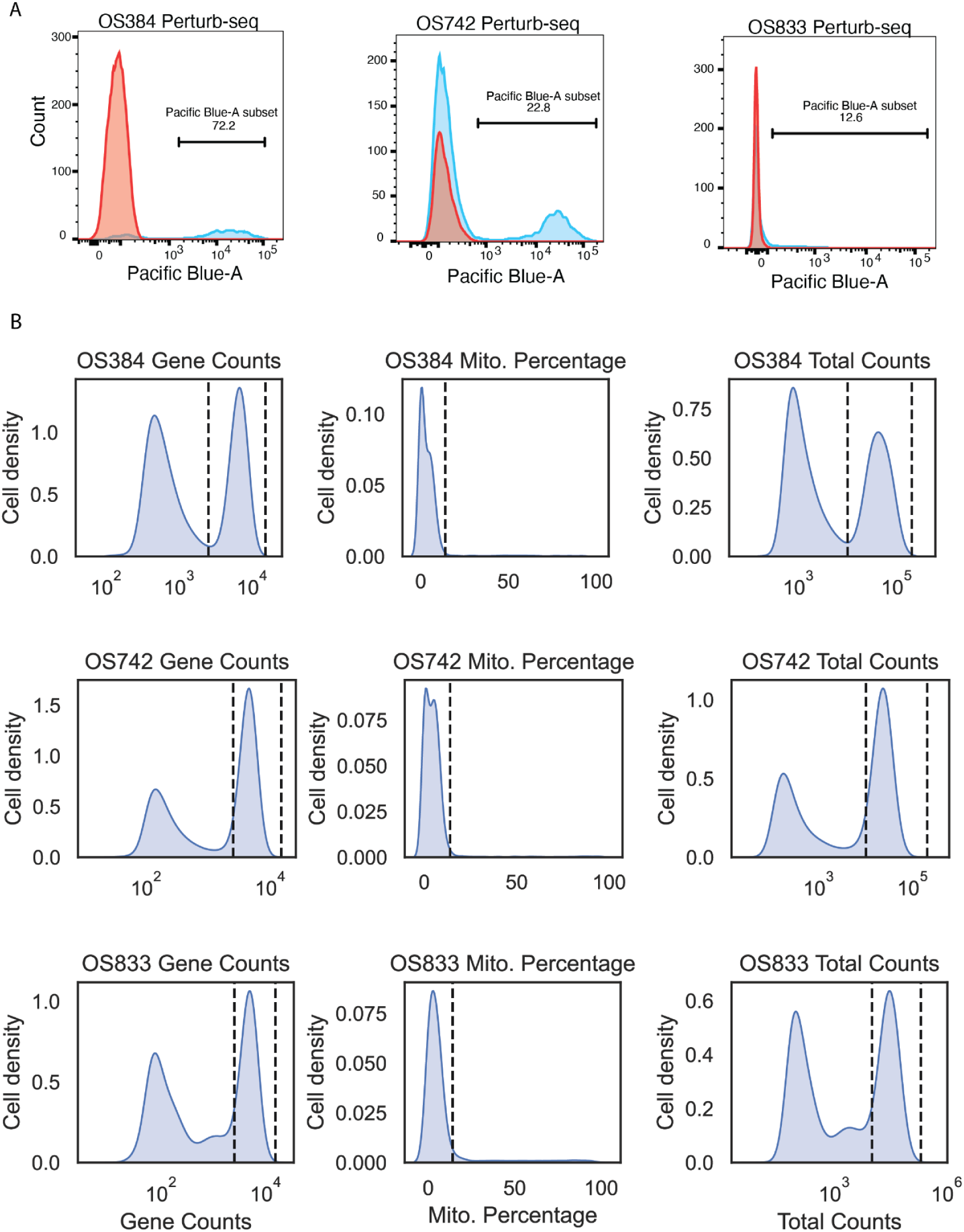
Perturb-seq sorting and QC. **(A)** OS PDX-derived cell lines were transduced with dCas9/CRISPRi KRAB constructs expressing mCherry, sorted based on mCherry and subsequently transduced with dual guide constructs and sorted based on both BFP and mCherry. **(B)** Quality control plots for all 3 samples used in the perturb-seq experiment. Datasets for OS384, OS742 and OS833 were filtered based on gene counts, mitochondrial percentage and total counts.

**Supplementary Figure 8:**
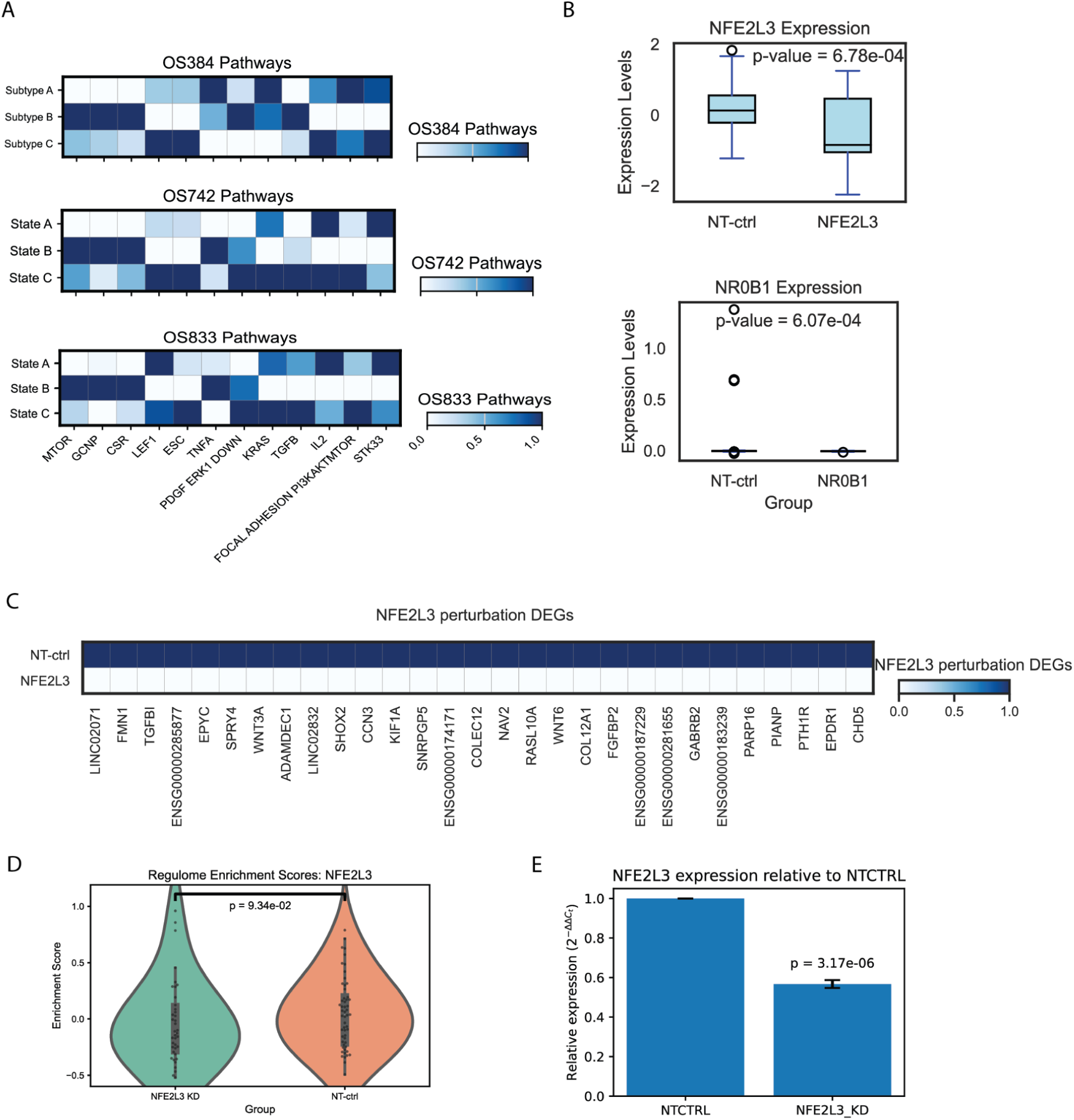
Perturb-seq QC and additional perturb-seq replicate. **(A)** Matrix plots representing quantified pathway enrichment in the different states. This analysis was performed on n=3 biological replicates. **(B)** Box plot comparing the expression levels of NFE2L3 between non-targeting control (NT-ctrl) and NFE2L3 perturbation groups. The box plot displays the median expression, interquartile range (light blue box), and whiskers representing the data range within 1.5 times the interquartile range. A Mann-Whitney U test was conducted to assess the statistical significance of the difference between the two groups, with the resulting p-value annotated on the plot (p-value = 6.78e-4). **(C)** The matrix plot displays differentially expressed genes (DEGs) following NFE2L3 perturbation in the OS833 sample compared to the non-targeting control (NT-ctrl). The heatmap is restricted to significant DEGs with a log_2_ fold change > 1 or <-1 and an adjusted p-value < 0.05. **(D)** Here, we are plotting the distribution of NFE2L3 regulome enrichment scores in the non-targeting control cells and the NFE2L3 KD cells. Statistical significance was determined using a Mann-Whitney U test. **(E)** Relative NFE2L3 expression in the NFE2L3 KD compared to the NT-CTRL cells. Significance was determined using the Welch’s t-test.

## Bibliography

1. Yabo YA, Niclou SP, Golebiewska A. Cancer cell heterogeneity and plasticity: A paradigm shift in glioblastoma. Neuro-Oncol. 2022;24(5):669–682. doi:10.1093/neuonc/noab269

2. Bergholz JS, Zhao JJ. How Compensatory Mechanisms and Adaptive Rewiring Have Shaped Our Understanding of Therapeutic Resistance in Cancer. Cancer Res. 2021;81(24):6074–6077. doi:10.1158/0008-5472.CAN-21-3605

3. Ottaviani G, Jaffe N. The Epidemiology of Osteosarcoma. In: Jaffe N, Bruland OS, Bielack S, eds. Pediatric and Adolescent Osteosarcoma. Cancer Treatment and Research. Springer US; 2010:3–13. doi:10.1007/978-1-4419-0284-9_1

4. Du X, Wei H, Zhang B, et al. Molecular mechanisms of osteosarcoma metastasis and possible treatment opportunities. Front Oncol. 2023;13. doi:10.3389/fonc.2023.1117867

5. Smeland S, Bielack SS, Whelan J, et al. Survival and prognosis with osteosarcoma: outcomes in more than 2000 patients in the EURAMOS-1 (European and American Osteosarcoma Study) cohort. Eur J Cancer. 2019;109:36–50. doi:10.1016/j.ejca.2018.11.027

6. Meltzer PS, Helman LJ. New Horizons in the Treatment of Osteosarcoma. N Engl J Med. 2021;385(22):2066–2076. doi:10.1056/NEJMra2103423

7. Shimizu T, Ishikawa T, Sugihara E, et al. c-MYC overexpression with loss of Ink4a/Arf transforms bone marrow stromal cells into osteosarcoma accompanied by loss of adipogenesis. Oncogene. 2010;29(42):5687–5699. doi:10.1038/onc.2010.312

8. Kamimoto K, Stringa B, Hoffmann CM, Jindal K, Solnica-Krezel L, Morris SA. Dissecting cell identity via network inference and in silico gene perturbation. Nature. 2023;614(7949):742-751. doi:10.1038/s41586-022-05688-9

9. Gong L, Yan Q, Zhang Y, Fang X, Liu B, Guan X. Cancer cell reprogramming: a promising therapy converting malignancy to benignity. Cancer Commun. 2019;39(1):48. doi:10.1186/s40880-019-0393-5

10. Rajan S, McAloney CA, Vetter TA, et al. Osteosarcoma Tumors Maintain Intra-Tumoral Heterogeneity, Even While Adapting to Environmental Pressures That Drive Clonal Selection.; 2021:2020.11.03.367342. doi:10.1101/2020.11.03.367342

11. Zhou Y, Yang D, Yang Q, et al. Single-cell RNA landscape of intratumoral heterogeneity and immunosuppressive microenvironment in advanced osteosarcoma. Nat Commun. 2020;11(1):6322. doi:10.1038/s41467-020-20059-6

12. Sheng G, Gao Y, Yang Y, Wu H. Osteosarcoma and Metastasis. Front Oncol. 2021;11. doi:10.3389/fonc.2021.780264

13. Itatani Y, Kawada K, Yamamoto T, Sakai Y. Resistance to Anti-Angiogenic Therapy in Cancer—Alterations to Anti-VEGF Pathway. Int J Mol Sci. 2018;19(4):1232. doi:10.3390/ijms19041232

14. Marchandet L, Lallier M, Charrier C, Baud’huin M, Ory B, Lamoureux F. Mechanisms of Resistance to Conventional Therapies for Osteosarcoma. Cancers. 2021;13(4):683. doi:10.3390/cancers13040683

15. Stewart E, Federico SM, Chen X, et al. Orthotopic patient-derived xenografts of paediatric solid tumours. Nature. 2017;549(7670):96-100. doi:10.1038/nature23647

16. Schott CR, Koehne AL, Sayles LC, et al. Osteosarcoma PDX-Derived Cell Line Models for Preclinical Drug Evaluation Demonstrate Metastasis Inhibition by Dinaciclib through a Genome-Targeted Approach. Clin Cancer Res. 2024;30(4):849–864. doi:10.1158/1078-0432.CCR-23-0873

17. Sayles LC, Breese MR, Koehne AL, et al. Genome-Informed Targeted Therapy for Osteosarcoma. Cancer Discov. 2019;9(1):46–63. doi:10.1158/2159-8290.CD-17-1152

18. Chen D, Zhao Z, Huang Z, et al. Super enhancer inhibitors suppress MYC driven transcriptional amplification and tumor progression in osteosarcoma. Bone Res. 2018;6:11. doi:10.1038/s41413-018-0009-8

19. Lopez R, Regier J, Cole MB, Jordan MI, Yosef N. Deep generative modeling for single-cell transcriptomics. Nat Methods. 2018;15(12):1053–1058. doi:10.1038/s41592-018-0229-2

20. Adamson B, Norman TM, Jost M, et al. A Multiplexed Single-Cell CRISPR Screening Platform Enables Systematic Dissection of the Unfolded Protein Response. Cell. 2016;167(7):1867–1882.e21. doi:10.1016/j.cell.2016.11.048

21. Li X, Dean DC, Cote GM, et al. Inhibition of ATR-Chk1 signaling blocks DNA double-strand-break repair and induces cytoplasmic vacuolization in metastatic osteosarcoma. Ther Adv Med Oncol. 2020;12:1758835920956900. doi:10.1177/1758835920956900

22. Hsu JY, Seligson ND, Hays JL, Miles WO, Chen JL. Clinical Utility of CDK4/6 Inhibitors in Sarcoma: Successes and Future Challenges. JCO Precis Oncol. 2022;(6):e2100211. doi:10.1200/PO.21.00211

23. He Y, Sun MM, Zhang GG, et al. Targeting PI3K/Akt signal transduction for cancer therapy. Signal Transduct Target Ther. 2021;6(1):425. doi:10.1038/s41392-021-00828-5

24. Ding L, Liu T, Qu Y, et al. lncRNA MELTF-AS1 facilitates osteosarcoma metastasis by modulating MMP14 expression. Mol Ther - Nucleic Acids. 2021;26:787–797. doi:10.1016/j.omtn.2021.08.022

25. Shijie L, Zhen P, Kang Q, Hua G, Qingcheng Y, Dongdong C. Deregulation of CLTC interacts with TFG, facilitating osteosarcoma via the TGF-beta and AKT/mTOR signaling pathways. Clin Transl Med. 2021;11(6):e377. doi:10.1002/ctm2.377

26. Lange M, Bergen V, Klein M, et al. CellRank for directed single-cell fate mapping. Nat Methods. 2022;19(2):159–170. doi:10.1038/s41592-021-01346-6

27. Saelens W, Cannoodt R, Todorov H, Saeys Y. A comparison of single-cell trajectory inference methods. Nat Biotechnol. 2019;37(5):547–554. doi:10.1038/s41587-019-0071-9

28. Mao M, Yuan Q, Xia X, Cui Y, Chen M, Yang W. Integrative analysis defines DDIT3 amplification as a correlative and essential factor for glioma malignancy. Am J Cancer Res. 2023;13(11):5418–5430.

29. Matherne MG, Phillips ES, Embrey SJ, Burke CM, Machado HL. Emerging functions of C/EBPβ in breast cancer. Front Oncol. 2023;13:1111522. doi:10.3389/fonc.2023.1111522

30. Dixit A, Parnas O, Li B, et al. Perturb-Seq: Dissecting Molecular Circuits with Scalable Single-Cell RNA Profiling of Pooled Genetic Screens. Cell. 2016;167(7):1853–1866.e17. doi:10.1016/j.cell.2016.11.038

31. Replogle JM, Norman TM, Xu A, et al. Combinatorial single-cell CRISPR screens by direct guide RNA capture and targeted sequencing. Nat Biotechnol. 2020;38(8):954–961. doi:10.1038/s41587-020-0470-y

32. Replogle JM, Bonnar JL, Pogson AN, et al. Maximizing CRISPRi efficacy and accessibility with dual-sgRNA libraries and optimal effectors. eLife. 11:e81856. doi:10.7554/eLife.81856

33. Tan X lan, Wang Z, Liao S, et al. NR0B1 augments sorafenib resistance in hepatocellular carcinoma through promoting autophagy and inhibiting apoptosis. Cancer Sci. 2024;115(2):465–476. doi:10.1111/cas.16029

34. Zhang XY, Zhang H, Hu SJ, et al. NR0B1 suppresses ferroptosis through upregulation of NRF2/c-JUN-CBS signaling pathway in lung cancer cells.

35. Chen T, Xu B, Chen H, et al. Transcription factor NFE2L3 promotes the proliferation of esophageal squamous cell carcinoma cells and causes radiotherapy resistance by regulating IL-6. Comput Methods Programs Biomed. 2022;226:107102. doi:10.1016/j.cmpb.2022.107102

36. Zhang M, Zhang YY, Chen Y, Wang J, Wang Q, Lu H. TGF-β Signaling and Resistance to Cancer Therapy. Front Cell Dev Biol. 2021;9:786728. doi:10.3389/fcell.2021.786728

37. Wang T, Wang D, Zhang L, et al. The TGFβ-miR-499a-SHKBP1 pathway induces resistance to EGFR inhibitors in osteosarcoma cancer stem cell-like cells. J Exp Clin Cancer Res CR. 2019;38:226. doi:10.1186/s13046-019-1195-y

38. Hattinger CM, Patrizio MP, Fantoni L, Casotti C, Riganti C, Serra M. Drug Resistance in Osteosarcoma: Emerging Biomarkers, Therapeutic Targets and Treatment Strategies. Cancers. 2021;13(12):2878. doi:10.3390/cancers13122878

39. Odri GA, Tchicaya-Bouanga J, Yoon DJY, Modrowski D. Metastatic Progression of Osteosarcomas: A Review of Current Knowledge of Environmental versus Oncogenic Drivers. Cancers. 2022;14(2):360. doi:10.3390/cancers14020360

40. Prudowsky ZD, Yustein JT. Recent Insights into Therapy Resistance in Osteosarcoma. Cancers. 2020;13(1):83. doi:10.3390/cancers13010083

41. Rajan S, Franz EM, McAloney CA, et al. Osteosarcoma tumors maintain intra-tumoral transcriptional heterogeneity during bone and lung colonization. BMC Biol. 2023;21(1):98. doi:10.1186/s12915-023-01593-3

42. Goyal Y, Dardani IP, Busch GT, et al. Pre-determined diversity in resistant fates emerges from homogenous cells after anti-cancer drug treatment. Published online December 9, 2021:2021.12.08.471833. doi:10.1101/2021.12.08.471833

43. Georgakopoulos-Soares I, Chartoumpekis DV, Kyriazopoulou V, Zaravinos A. EMT Factors and Metabolic Pathways in Cancer. Front Oncol. 2020;10:499. doi:10.3389/fonc.2020.00499

44. Liu S, Ren J, ten Dijke P. Targeting TGFβ signal transduction for cancer therapy. Signal Transduct Target Ther. 2021;6(1):1–20. doi:10.1038/s41392-020-00436-9

45. Qian J, Huang C, Zhu Z, et al. NFE2L3 promotes tumor progression and predicts a poor prognosis of bladder cancer. Carcinogenesis. 2022;43(5):457–468. doi:10.1093/carcin/bgac006

46. Ren Y, Yang J, Ding Z, et al. NFE2L3 drives hepatocellular carcinoma cell proliferation by regulating the proteasome-dependent degradation of ISGylated p53. Cancer Sci. 2023;114(9):3523–3536. doi:10.1111/cas.15887

47. Köksal H, Müller E, Inderberg EM, Bruland Ø, Wälchli S. Treating osteosarcoma with CAR T cells. Scand J Immunol. 2019;89(3):e12741. doi:10.1111/sji.12741

48. Yeo GHT, Lin L, Qi CY, Cha M, Gifford DK, Sherwood RI. A Multiplexed Barcodelet Single-Cell RNA-Seq Approach Elucidates Combinatorial Signaling Pathways that Drive ESC Differentiation. Cell Stem Cell. 2020;26(6):938–950.e6. doi:10.1016/j.stem.2020.04.020

49. Lotfollahi M, Susmelj AK, Donno CD, et al. Learning interpretable cellular responses to complex perturbations in high-throughput screens. bioRxiv. Published online May 18, 2021:2021.04.14.439903. doi:10.1101/2021.04.14.439903

50. Roohani Y, Huang K, Leskovec J. Predicting transcriptional outcomes of novel multigene perturbations with GEARS. Nat Biotechnol. Published online August 17, 2023:1–9. doi:10.1038/s41587-023-01905-6

51. Hsiung CCS, Wilson CM, Sambold NA, et al. Engineered CRISPR-Cas12a for higher-order combinatorial chromatin perturbations. Nat Biotechnol. Published online May 17, 2024:1–15. doi:10.1038/s41587-024-02224-0

52. Schott CR, Koehne AL, Sayles LC, et al. Development and characterization of new patient-derived xenograft (PDX) models of osteosarcoma with distinct metastatic capacities. Published online January 20, 2023:2023.01.19.524562. doi:10.1101/2023.01.19.524562

53. La Manno G, Soldatov R, Zeisel A, et al. RNA velocity of single cells. Nature. 2018;560(7719):494-498. doi:10.1038/s41586-018-0414-6

54. Tirosh I, Izar B, Prakadan SM, et al. Dissecting the multicellular ecosystem of metastatic melanoma by single-cell RNA-seq. Science. 2016;352(6282):189-196. doi:10.1126/science.aad0501

55. Bergen V, Lange M, Peidli S, Wolf FA, Theis FJ. Generalizing RNA velocity to transient cell states through dynamical modeling. Nat Biotechnol. 2020;38(12):1408–1414. doi:10.1038/s41587-020-0591-3

56. Bergen V, Lange M, Peidli S, Wolf FA, Theis FJ. Generalizing RNA velocity to transient cell states through dynamical modeling. Nat Biotechnol. 2020;38(12):1408–1414. doi:10.1038/s41587-020-0591-3

57. Adamson B, Norman TM, Jost M, et al. A Multiplexed Single-Cell CRISPR Screening Platform Enables Systematic Dissection of the Unfolded Protein Response. Cell. 2016;167(7):1867–1882.e21. doi:10.1016/j.cell.2016.11.048

58. Lange M, Bergen V, Klein M, et al. CellRank for directed single-cell fate mapping. Nat Methods. 2022;19(2):159–170. doi:10.1038/s41592-021-01346-6

